# The archaeal triphosphate tunnel metalloenzyme *Sa*TTM defines structural determinants for the diverse activities in the CYTH protein family

**DOI:** 10.1101/2021.03.18.435988

**Authors:** Marian S. Vogt, Roi R. Ngouoko Nguepbeu, Michael K. F. Mohr, Sonja-Verena Albers, Lars-Oliver Essen, Ankan Banerjee

## Abstract

CYTH is a large protein superfamily that is conserved in all three domains of life with its unique triphosphate tunnel metalloenzyme (TTM) fold. Besides phosphatase functions, e.g. as RNA triphosphatase, inorganic polyphosphatase or thiamine triphosphatase, some CYTH orthologs cyclize nucleotide triphosphates to 3’,5’-cyclic nucleotides. So far, archaeal CYTH proteins are annotated as adenylyl cyclases although experimental evidence is lacking. To address this gap, we characterized a CYTH ortholog, *Sa*TTM, from the crenarchaeote *Sulfolobus acidocaldarius*. Our initial *in silico* studies suggested a close relationship between archaeal CYTH enzymes and class IV adenylyl cyclases compared to the other CYTH-subclasses, but biochemical data showed no cyclic nucleotide production. Instead, our structural and functional analyses show a classical TTM behavior. The Ca^2+^-inhibited Michaelis complex indicates a two-metal ion reaction mechanism analogous to other TTMs. Different co-crystal structures of *Sa*TTM further reveal conformational dynamics in *Sa*TTM, let us to assume feedback inhibition in TTMs due to tunnel closure in the product state. Combining our structural insights with sequence-similarity network based *in silico* analysis, we further set out a firm molecular basis for distinguishing CYTH orthologs with phosphatase activities from class IV adenylyl cyclases.

**Major highlights:**

- CyaB-like class IV adenylyl cyclase homologs in archaea are triphosphatases.
- The co-crystal structure of *Sa*TTM in sulfate and triphosphate bound state revealed conformational transition of the TTM tunnel during catalysis.
- Atomic insights into TTM inhibition by calcium and pyrophosphate.
- *In silico* and structure-function analysis revealed the molecular determinant for functional diversification among CYTH proteins.

## Introduction

Bacterial <u>Cy</u>aB-like class IV adenylyl cyclases (AC-IV) and mammalian thiamine triphosphatases (<u>Th</u>TPase) are founding members of the *CYTH-like domain superfamily* (1). CYTH enzymes are found in all three domains of life and can be traced back to the last common ancestor, which makes them to appealing targets for studying protein evolution (1). Beside CyaB and ThTPases, the *CYTH-like domain superfamily* comprises inorganic triphosphatases (2, 3), which together are wrapped up in a *CYTH domain* subfamily, and Cet1-like RNA-triphosphatases that have their own subfamily of *mRNA triphosphatase Cet1-like* enzymes within the superfamily (4).

The characteristic structural feature of CYTH enzymes is an 8-stranded β-barrel tunnel architecture, termed triphosphate tunnel metalloenzyme (TTM) fold, which was first found in the crystal structure of the fungal Cet1 RNA triphosphatase (4, 5). The hallmark of CYTH enzymes is an N-terminal *ExExK* motif that is involved in metal ion coordination and substrate binding (3). The opposing acidic and basic patches shape the substrate binding tunnel. While the overall coordination of the triphosphate itself is very similar among CYTH enzymes, the directionality of asymmetric organic substrates such as triphosphate nucleotides or thiamine triphosphate within the tunnel in fact determines catalytic specificity (3). For triphosphatases and ThTPases, the organic moiety points towards a C-terminal helix, termed plug helix (3, 6). Such enzymes catalyze the hydrolysis of the triphosphate by a two-metal ion mechanism using metal 1 to establish the catalytically competent binding of the triphosphate moiety and metal 2 to position and activate a water for nucleophilic attack onto the γ-phosphate moiety (3). In the case of triphosphatases, a pyrophosphate and an orthophosphate are produced by hydrolysis of the β—γ phosphoanhydride bond. The situation for AC-IV enzymes in the CYTH superfamily is different. Here, the directionality of the organic moiety binding within the tunnel fold is inverted causing a reversed placement of the α- and γ-phosphates within the binding tunnel (7). By comparing AC-IV with already described AC-classes, a general two-metal ion mechanism was suggested for the cyclization of ATP to 3’,5’-cAMP using the second metal to lower the activation energy (7–10).

The founding members of the CYTH superfamily are involved either in mammalian thiamine triphosphate metabolism or bacterial signal transduction via 3’,5’-cyclic nucleotides. Some members of the CYTH superfamily are parts of multi-domain proteins by being fused to HD hydrolase, exopolyphosphatase, nucleotidyl kinase, or CHAD domains (1). It was postulated that CYTH protein might play a central role at the interface between nucleotide and phosphate metabolism (1, 11). Archaeal representatives encode at least one copy of a CYTH ortholog in their genomes, which are generally annotated as adenylyl cyclases. The presence of 3’,5’-cAMP is described for a few archaeal species indicating a role in starvation and cell cycle (12, 13). In search for a potential nucleotidyl cyclase in the crenarchaeote *Sulfolobus acidocaldarius*, we identified *Saci_0718*, a CYTH superfamily protein, annotated as CyaB-like class IV adenylyl cyclase. Although the substrate binding mode and the hydrolysis mechanisms differ between triphosphatases and CyaB (3), at the sequence level, *Saci_*0718 shares ∼29% identity with the *Yersinia pestis* cyclase CyaB and with the inorganic triphosphatase from *Clostridium thermocellum*.

Here, we have performed a structure-function analysis for the only ortholog of a CYTH enzyme in *Sulfolobus acidocaldarius, Sa*TTM. *Sa*TTM is a triphosphatase. Its structures in sulfate and substrate (triphosphate and ATP) bound states revealed conformational dynamics during catalysis. We trapped the Michaelis-complex of the enzyme in its calcium-inhibited state that established TTM phosphatases follow a two-metal ion mechanism. The product-bound (PPi) co-crystal structure set the molecular basis of feedback inhibition in TTM. Our structure-function analysis identified the important role of ordered water molecules for substrate binding. Our in-depth sequence similarity network separated CYTH superfamily into various clusters according to their function. Taken together, a comparative structural and bioinformatic analysis of the CYTH proteins led us to identify motifs that represent molecular determinants underpinning the different functions as adenylyl cyclase and phosphatase within the CYTH enzymes.

## Results

### Archaeal CYTH proteins are functionally diverged from CyaB-like class IV adenylyl cyclases

To date, the *CYTH-like domain superfamily* (IPR033469) comprises 41000 enzymes from all three domains of life divided in two subfamilies, the smaller *mRNA triphosphatase Cet1-like* (IPR004206) and the bigger subfamily of *CYTH domain* proteins (IPR023577) including adenylyl cyclases, ThTPases, and inorganic tri- or polyphosphatases (also known as triphosphatase tunnel metalloenzymes or TTMs). The discrimination between *Cet1-like* and *CYTH* enzymes is also consistent within the Pfam database (*mRNA_triPase:* PF02940 and *CYTH:* PF01928). Hence, we aimed to obtain a detailed understanding of the *CYTH* family by a detailed bioanalytical analysis combined with structural insights, as the functional annotations of *CYTH* enzymes is still vague and misleading among TTMs and adenylyl cyclases.

In order to restrict our studies on an unambiguous assignment, we chose the Pfam family *CYTH* (PF01928) as the least common denominator for our analysis. As stated above, this protein family contains enzymes with known functions as TTMs, AC-IVs, and ThTPases. We therefore performed a systematic sequence similarity network (SSN) analysis of the CYTH superfamily PF01928 using EFI-EST (14). The network generated with an E-value of 10^−20^ against the *CYTH*-domain boundaries for 38246 sequences, clustered to 4128 nodes with pairwise sequence identities of >40% within the nodes, which are connected by a total of 121791 edges. This SSN analysis covering all members of the *CYTH* enzymes of PF01928 showed ten major distinct clusters (Fig. 1). Among these clusters, three of them (cluster 1, 4 and 7) comprise members adopting a multi-domain architecture, while the length histograms for the remaining seven clusters are in the expected range of ∼160-200 amino acids for single domain proteins (Fig. S7A). For example, cluster 1 contains YgiF from *Escherichia coli*, which possesses a two-domain architecture with an N-terminal CYTH and a C-terminal CHAD domain (PF05235) (3). Similar modular architectures including the CHAD-domain can be found in cluster 4, which does not contain any structurally described homologs and whose members occur mostly in actinobacteria. The additional domain in cluster 7, whose members are exclusively found in *Viridiplantae* and *Amoebozoa*, is a phosphoribulokinase domain (PF00485).

**Figure 1.**
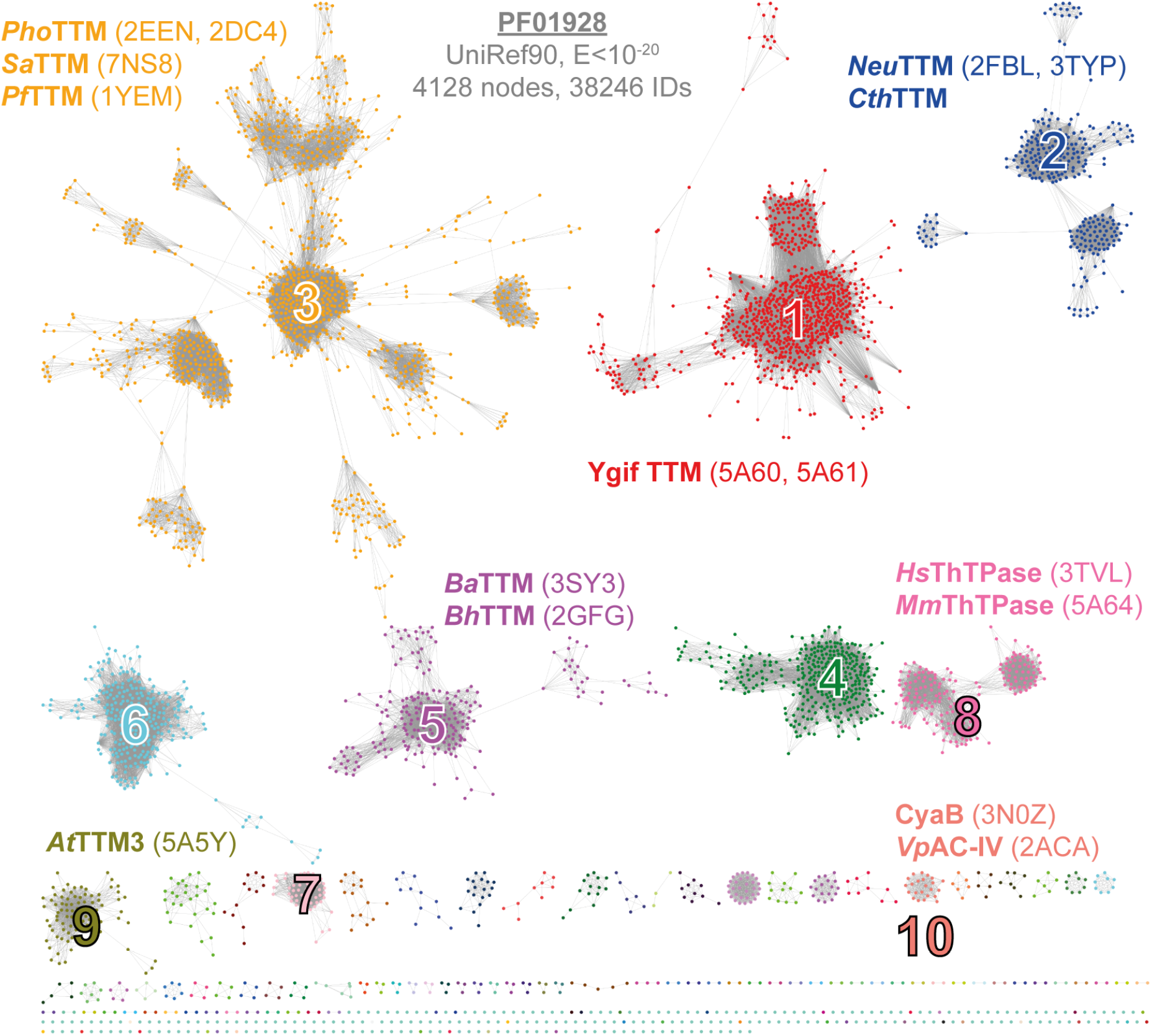
Sequence similarity network of CYTH proteins. The CYTH family PF01928 is presented as a sequence-similarity network, generated with the EFI-EST using the domain boundaries and a final alignment score of 10^−20^ (14). Each node represents sequences with >40% identity and the final network was subjected to the cluster analysis implemented by the EFI-EST. Accordingly, the ten largest clusters were subjected for further analysis, including all currently described CYTH enzymes (see Table S1). Known TTMs are found in clusters 1-3, 5, and 9. ThTPase enzymes appear exclusively in cluster 8 and CyaB/AC-IV’s belong to cluster 10. Proteins from cluster 4 are unknown and cluster 7 members lack the hallmark motif ExExK. If available, PDB entries are given in parentheses. The network with E<10^−15^ and WEBLOGOs of clusters 1-10 are shown in Figure S7+8.

Interestingly, the SSN analysis discriminates the cyaB-like class IV adenylyl cyclase (7, 15) (cluster 10) from other triphosphatases. Cluster 10 is a relatively small cluster within the SSN by comprising only 141 orthologs of bacterial origin. Like AC-IV, the ThTPases form a separate cluster, cluster 8, that is distinct from other triphosphatases. The key molecular determinant of ThTPase was reported previously (3, 6). The enzymes described as inorganic triphosphatases are distributed among different clusters. *Neu*TTM and *Cth*TTM were found in cluster 2, TTM3 from *Arabidopsis thaliana* is a member of cluster 9 (3, 16, 17). The two other described enzymes from *A. thaliana* (TTM1 and TTM2) are in cluster 7 and interestingly they utilize pyrophosphate and not triphosphate (18). *At*TTM1 and *At*TTM2 does not possess the TTM-signature motifs required for catalysis (Table S1). In cluster 5, unpublished structures annotated before as adenylyl cyclase from *Bacillus anthracis* and *B. halodurans* can be found. A member of this cluster from *Staphylococcus aureus* has further been described to be incompetent in cAMP formation and to be distinct from class IV adenylyl cylases (19).Cluster 3 contains orthologs of mostly archaeal origin from *Pyrococcus furiosus* and *P. horikoshii*, as well as *Saci_*0718 (Uniprot: *Q4JAT2*). The SSN analysis clearly indicating that the archaeal CYTH enzymes are functionally diverse from CyaB-like class IV adenylyl cyclases.

### *Sulfolobus acidocaldarius CYTH* enzyme is a triphosphatase tunnel metalloenzyme

To gain in depth biochemical and structural insights into this protein, we heterologously overproduced a C-terminal His_6_-tagged variant of *Saci*_0718 in *Escherichia coli*. The recombinant protein was purified to homogeneity using an initial heat step followed by IMAC and subsequent size exclusion chromatography as a polishing step. The enzyme formed a dimer in solution corresponding to 56 kDa compared to standards (Fig. S1). Dimerization behavior is typical among other CYTH proteins (11). The first ambiguity to be resolved was determination of its substrate preference and the nature of its hydrolysis products. Initially, we tested its activity against ATP using Mg^2+^ as cofactor at 75 °C and the reaction product analyzed by HPLC (Fig. S2A). This experiment clearly showed that the enzyme is capable to hydrolyze ATP to ADP and Pi without any formation of cAMP. *Saci*_0718 showed no base specificity during its phosphohydrolase reaction and is able to utilize both ATP or GTP with similar efficiency (Fig. 2A). As the reaction products are diphosphate nucleosides and not (c)AMP/GMP, we used non-hydrolysable analogs of ATP, AMPCPP and AMPPCP, to identify the nucleophilic attack site (Fig. S2B). It shows that Saci_0718 acts on the terminal phosphate of ATP, as the enzyme was active on AMPCPP but no phosphate release was observed for AMPPCP.

**Figure 2.**
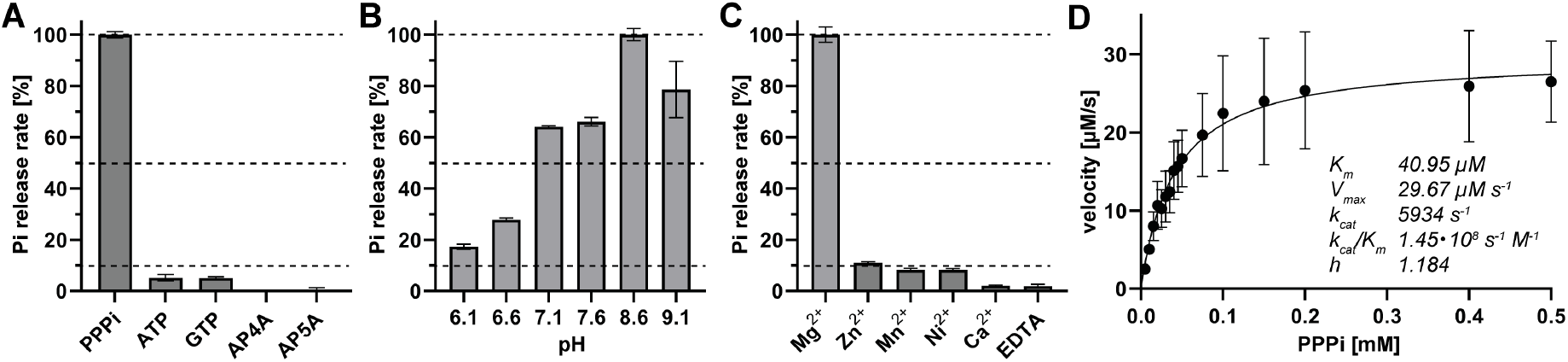
The main substrate of *Sa*TTM is PPPi and not nucleotides. **A** For substrate specificity, 5 nM *Sa*TTM were incubated with 2 mM MgCl2 and 0.1 mM substrate at pH 8.6 and 75 °C for 10 min. **B** The pH optimum of *Sa*TTM peaks at pH 8.6 within the tested range of 6.1 to 9.1, using the conditions from **A. C** Different metal ions (Mg^2+^, Zn^2+^, Mn^2+^, Ni^2+^, and Ca^2+^, EDTA as control, 2 mM each) were examined to show their effect on *Sa*TTM activity, using the conditions from **A** and an incubation time of 5 min. **D** v/S diagram of *Sa*TTM using 5 nM enzyme, 0.005-0.5 mM PPPi as substrate at pH 8.6 and 75 °C. The kinetic parameters are noted.

Next, we used triphosphate (PPPi) as substrate. Under the conditions outlined before *Saci_0718* is ∼19 times more active compared to NTP as substrate. The result is consistent with the previous reports of tunnel metalloenzymes (2, 3). We also observed that the enzyme is able to hydrolyze higher polyphosphates like tetraphosphate with final formation of pyrophosphate and phosphate (Fig. S3). As triphosphate seems to be the preferred substrate, we named *Saci_0718 Sa*TTM (*<u>S</u>. <u>a</u>cidocaldarius* triphosphate tunnel <u>m</u>etalloenzyme). The optimal assay conditions were observed at a basic pH 8.6 at 75 °C using Mg^2+^ as a cofactor, while Zn^2+^, Mn^2+^, and Ni^2+^ could be used as weaker cofactor alternatives. Similar to the metal ion chelator EDTA, calcium exerted an inhibitory effect on phosphatase activity (Fig. 2B-C). At optimal conditions, kinetic measurement revealed a classical Michaelis-Menten behavior with a *K*_*m*_ of 40.95 µM and a *V*_*max*_ of 29.67 µM s^−1^. This shows a highly efficient catalysis performed by *Sa*TTM of 1.45×10^8^ s^−1^ M^−1^ close to the diffusion limit and a turnover rate (5934 s^−1^), which is in the range of untagged *Neu*TTM (7900 s^−1^) (16).

### Crystal structure of the *Sa*TTM dimer

For structural information, we crystallized *Sa*TTM in different catalytically relevant complexes. Initial attempts of producing diffraction quality crystals using apo enzyme were unsuccessful. Only by adding ATP as ligand, diffraction quality crystals were obtained, indicating that *Sa*TTM is highly flexible in solution and stabilized upon substrate binding. Orthorhombic crystals of *Sa*TTM in its different catalytic states were obtained after a week at 4°C that diffracted to 1.75-2.3 Å resolution. For structure determination, *Pyrococcus horikoshii* OT3 *PH1819* (PDB entry 2EEN) was used as search model in molecular replacement. *Sa*TTM was crystallized as a relaxed homodimer (Fig. 3A), bound with two sulfate anions instead of the ATP that was included in co-crystallization. Of note, a solution structure of mouse thiamine triphosphatase (PDB 2JMU) reported a similarly relaxed conformation. The open state of *Sa*TTM was crystallized in space group *P*4_1_2_1_2 with one monomer per asymmetric unit. The polypeptide is defined by electron density for R6-A176, but interrupted for P44-R49 and Q73-R80 by missing density. The general topology of *Sa*TTM is β1-α1-β2-β3-β4-β5-α2-β6-β7-β8-α3-β9-α4. The dimerization of *Sa*TTM was also found in the crystal structure formed by two symmetry mates. PDBePISA revealed that the dimer interface covered 875 Å^2^ with a CSS score of 0.32 and accounts for 8.5% of the total solvent-accessible area of one protomer (20). The dimerization is mainly established by hydrophobic interactions of residues from β3, β4, β5, and α2 and salt bridges between K99 from one protomer with E81 from the other protomer. The C-terminal α4 plug helix blocks the tunnel entry from one side. It was previously reported that this plug helix plays a major role in determining substrate preference (2).

**Figure 3.**
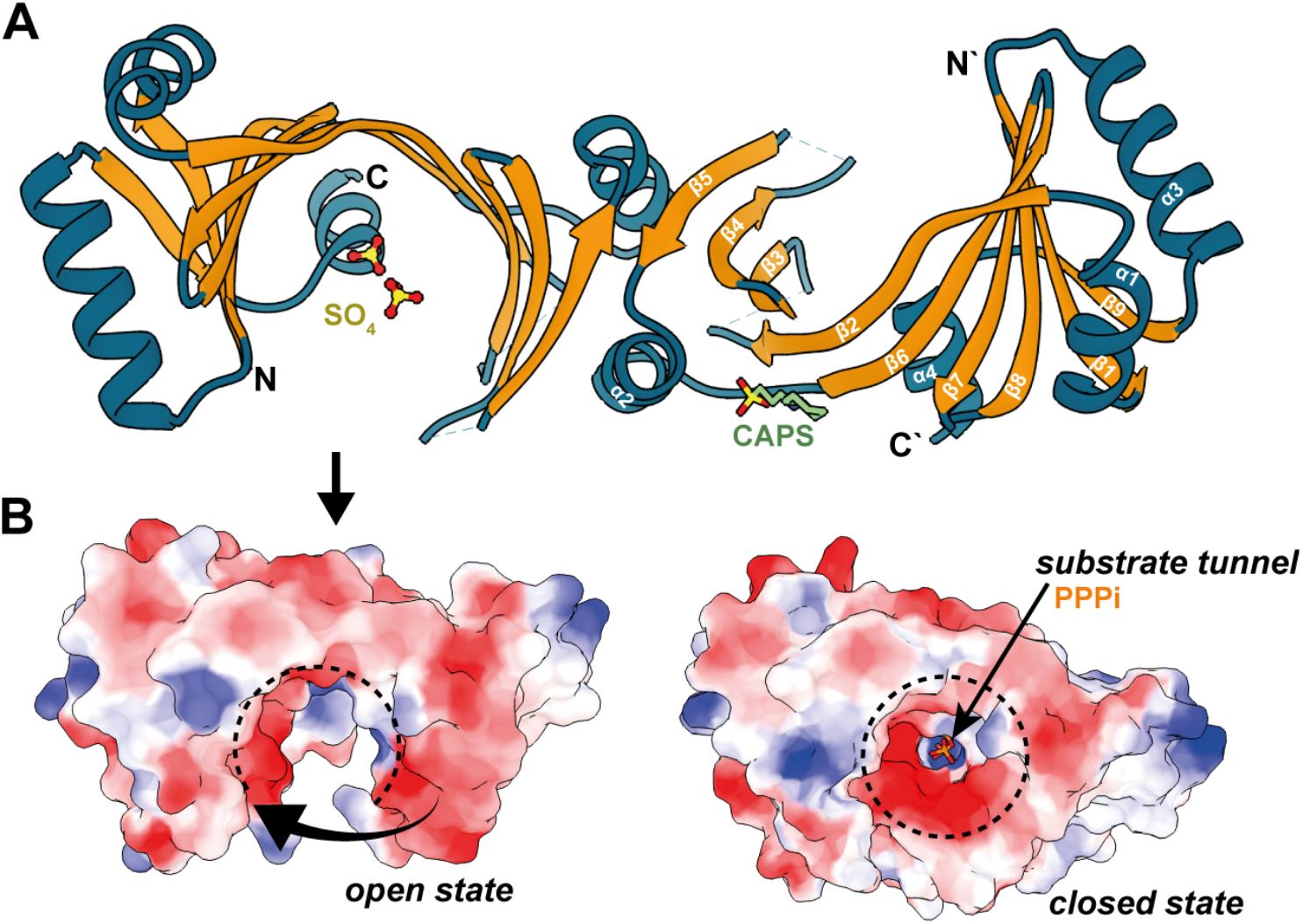
The homodimer of SaTTM has an open and closed state. **A** The relaxed or open homodimer of SaTTM is shown as cartoon with the beta barrel being highlighted in orange and the rest being colored in petrol. The secondary structure elements are indicated on the right protomer. The cocrystallized sulfates and CAPS molecules (sticks with green carbons, red oxygens, and yellow sulfurs) are shown in one of the protomers for clarity, but are present in both, as the dimer has been generated by the symmetry mate due to just one molecule per AU. N- and C-termini of the protomers are indicated, the dashed lines indicate missing electron density for residues P44—R49 and Q73-R80. **B** The left panel shows the above orientation of the open states’ surface as indicated by the arrow using the electrostatic coloring from negative (red) to positive (blue). The arrow indicates the closing movement upon ligand binding, resulting in the closed conformation shown on the right with the PPPi (orange and red sticks) substrate in orange bound within the tunnel. The dashed (open) circles indicate the conformation switch.

### The catalytic ensemble of *Sa*TTM

Next, we aimed to trap the Michaelis complex of *Sa*TTM, for which we co-crystallized the enzyme together with PPPi or ATP, and the respective cofactors Mg^2+^ or Ca^2+^ at 4 °C. Substrate-bound states of *Sa*TTM crystallized in a closed tunnel conformation, highlighting the structural rearrangement upon ligand binding (Fig. 3B). This supports the previous notion that CYTH enzymes are highly flexible and can breathe during their catalytic cycle (6, 17). In all states but the open state, a closed tunnel conformation including the missing densities of P44-R49 and Q73-R80 from the open state is observed, providing a continuous density map throughout the polypeptide.

As shown for *At*TTM3 (3), we observed electron density in the PPPi.Mg^2+^ state for just one metal ion octahedrally coordinated by PPPi, E7, and E135 (Fig. 4A and S4A), while the second ion necessary for water activation was missing (Fig. S5). When using a complete PPPi molecule during refinements, we found higher B-factors for the γ-phosphate. Therefore, the molecule was split into PPi and γ-Pi and occupancies were separately refined. This revealed that only 76% of the substrate binding site were occupied with γ-phosphates, suggesting a slow turnover of the triphosphate *in crystallo*.

**Figure 4.**
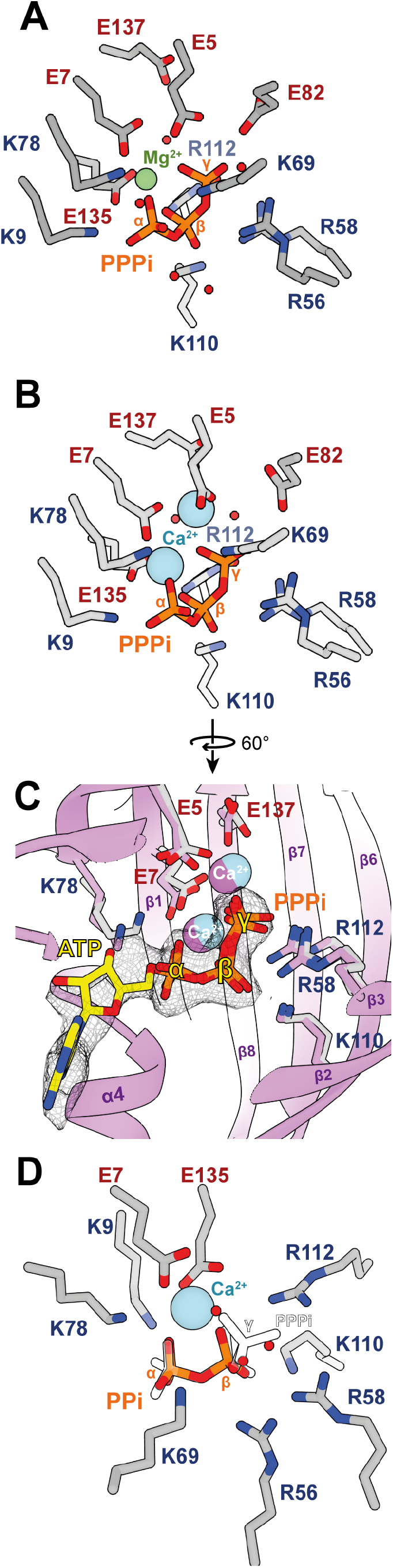
Catalytic states of SaTTM. **A** Residues (grey) in the binding tunnel of SaTTM are shown coordinating its substrate PPPi (orange sticks) and a magnesium ion (green sphere) at metal binding site 1. **B** Same orientation as in A of residues (grey) binding PPPi and two calcium ions (blue sphere) is shown. **C** For identification of triphosphate orientation, SaTTM is cocrystallized with ATP (yellow) and two calcium ions in the context of secondary structure elements (as indicated) and residues in purple. The superposition with the PPPi-calcium state from B is shown with the indicated residues to assign the α - and γ-positions of the symmetric substrate in A, B, and D. **D** The product bound state of SaTTM in complex with PPi and calcium is shown with the same coloring scheme as in B. The PPPi is shown with contours only, identifying the α - and β-positions and further show that the three indicated waters (red spheres) localize at the γ-phosphate oxygen positions.

The PPPi.Ca^2+^ state (Fig. 4B) revealed not only metal ion 1, but also the water-activating position of metal ion 2 site present in the electron density. Two Ca^2+^ ions are coordinated by PPPi, E5, E7, and E137 (Fig. S4B), thereby positioning a water molecule in an inline-attack position 3.2 Å away from the γ-phosphate phosphorous as observed for the *E. coli* YgiF.PPPi states (PDB: 5A60, 5A61). Here, no differences of γ-phosphate B-factors and occupancies were observed, indicating no hydrolysis during crystallization, which is in line with activity tests in the presence of Ca^2+^ (Fig. 2C). As *Sa*TTM catalyzes asymmetrical cleavage of a symmetric substrate it is difficult to judge the directionality of the substrate, a prerequisite for firm mechanistic conclusions. Hence, we solved the ATP-bound form of *Sa*TTM in the presence of Ca^2+^ ions. The *Sa*TTM.ATP state revealed the adenosine base on the side of the plug helix α4, defining the orientation of the triphosphate moiety. It also showed that the catalytic mechanism of *Sa*TTM follows a classical inorganic triphosphatase, but not an AC-IV, providing the structural base of TTM function (Fig. 4C). Upon the structural and biochemical confirmation of *Sa*TTM’s inability to use calcium as reaction-competent cofactor we wondered how Ca^2+^ competes with Mg^2+^. Hence, we performed a titration experiment with increasing concentrations of calcium on a magnesium-catalyzed reaction (Fig. S6A). As expected, we observed at equimolar concentrations a highly reduced activity of *Sa*TTM using PPPi as substrate, and almost no turnover with 5-fold excess of the inhibiting metal ion, indicating that calcium is a potent inhibitor of TTM.

For comparison with the product state, we determined the crystal structure of *Sa*TTM in complex with pyrophosphate (Fig. 4D). As the Michaelis complex, the post catalytic state appeared to have the same closed tunnel conformation (Fig. S6). By using the superimposed triphosphate state, the two phosphates can be assigned to the α - and β-positions complexing the calcium ion at metal 1 position (Fig. 4D). Three water molecules occupy the positions corresponding to the oxygens from the substrate’s γ-phosphate. The product bound state clearly suggests an inhibitory effect of pyrophosphate on TTM activity and to confirm this notion we performed another titration-inhibition experiment (Fig. S6B). The activity of *Sa*TTM was rapidly decreased and already halved when using a ten-thousandth of the substrate concentration. While this effect plateaus until equimolar concentrations, the inhibition upon tenfold excess is amplified and with 100-fold excess the enzyme activity was completely abolished. The tunnel closure upon product binding and the enzyme-titration indicates a feedback inhibition mechanism in TTMs.

### Role of ordered water molecule in substrate binding and catalysis

Interestingly, during SSN analysis we found at a less stringent alignment score of 10^−15^ that CYTH enzyme orthologs from cluster 3 (including *Sa*TTM) and cluster 10 (including CyaB) belong still to the same cluster. This suggests generally higher sequence and structural similarity between these CyaB and SaTTM-like enzymes compared to other TTM subfamilies, while the triphosphatase-like catalytic mechanism of *Sa*TTM is apparently shared with CYTH enzyme orthologs already well separated at less stringent thresholds. To find a structural rationalization for this phenomenon, we compared the substrate bound states of *Yp*CyaB, *At*TTM, and *Sa*TTM (Fig. 5A-B). We identified two water molecules, which are apparently prerequisites for substrate binding, as they occur in all known structures of the CYTH superfamily. These waters are located oppositely to the metal ions and directly hydrogen-bonded to the α - and β-phosphate, hence termed as α - and β-water (Fig. 5A). While β-water in *Sa*TTM and *Yp*CyaB made a bidentate contact with the DxY motif, this interaction is absent in *At*TTM due to a slightly moved water and an NxF motif. Here, the asparagine is 4 Å away and the phenylalanine lacks the ability to establish a hydrogen bond. This β-water establishes a hydrogen bond with R52 on β3, explaining the well-ordered β-water in *At*TTM (Fig. 5B). The deviations in substrate coordination can also be observed on the metal-ion binding side. While *Sa*TTM and *Yp*CyaB share the same arrangement by shaping the binding pocket in close proximity to the α-phosphate using a combination of two conserved aromatic residues (Y168/173 and F133/134), the situation in *At*TTM is different. Here, a salt bridge ensemble of E167 and K200 is used for direct electrostatic interaction with the γ-phosphate. As we observed this highly similar binding mode of α - and β-phosphates in *Sa*TTM and *Yp*CyaB, we wondered about the impact of these residues on enzyme activity. Hence, we mutated the DxY motif and the central tyrosine Y168, as well as E135 as a dead mutant control due to its central role in metal 1 binding (Fig. 5C). As hypothesized, the *Sa*TTM D38A and Y40A mutants showed reduced activity compared to the wild type. Intuitively, less impact on the catalysis was observed for the conservative D38N and Y40F mutants compared to the alanine mutants, as similar hydrogen bond formation is still possible in the asparagine variant, and in the phenylalanine variant we still have the aspartate for β-water coordination. The double mutant had an accumulating effect with less than 10% of wild type activity, strengthening the importance of the DxY motif in coordinating properly the water-substrate complex for catalysis. Interestingly, the Y168A mutant showed a twofold turnover rate compared to the wild type enzyme, while the corresponding K200A mutant in *At*TTM mutant showed reduced activity (3). These residues are part of the plug helix, a structural element that plays a critical role in substrate preference and enzyme activity (2).

**Figure 5.**
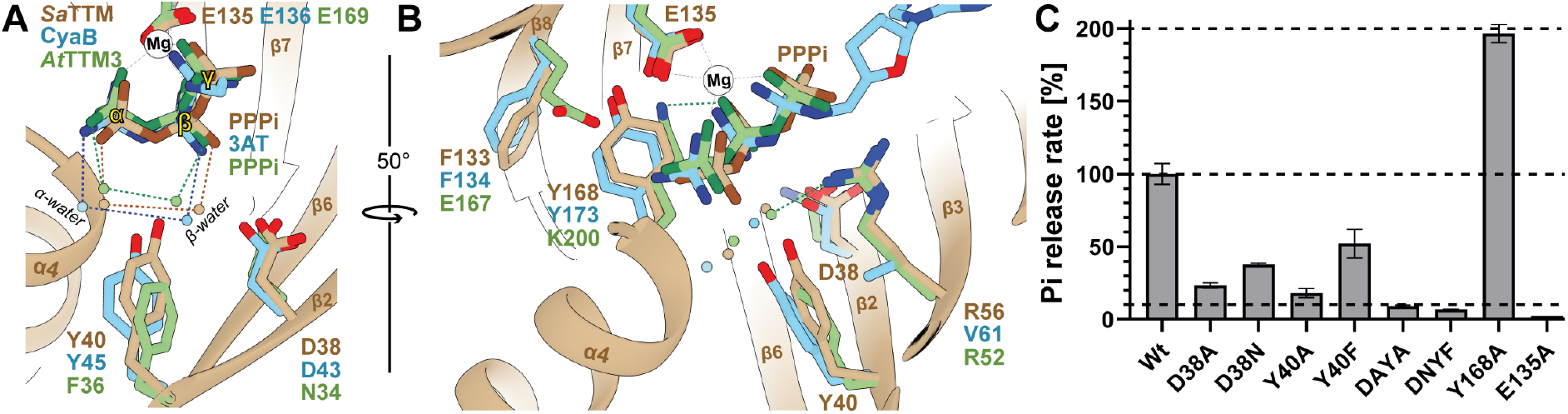
Different modes of substrate coordination among TTMs utilizing two conserved waters. **A+B** Superposition of substrate binding mode of CyaB (3N0Z, blue), AtTTM3 (5A5Y, green), and *Sa*TTM (7NS8, brown) is shown in the context of secondary structure elements of SaTTM from two different angles as indicated. The magnesium cofactor (white sphere) from SaTTM is shown. Although mechanistically different, the binding modes of CyaB and SaTTM are highly similar regarding the establishment of the substrate waters, while AtTTM3 differs strikingly. **B** PPPi hydrolysis activities of the indicated residues show the impact of correct water placement by the DxY-motif and a positive effect on activity of a loosened plug helix.

## Discussion

### Structural signature that defines versatile CYTH function

Decades of study on *CYTH* containing proteins from the protein family *PF01978* established the catalytic mechanisms. However, due to high sequence similarities between different isofunctional subclusters it proved hard to assign concrete functions to uncharacterized family members. We aimed to integrate our newly gained structural insights with our findings from the sequence similarity network and already known CYTH enzymes to establish a detailed understanding about their versatile enzyme activities. The directionality of substrate binding is a major mechanistic determinant on the obtained reaction product when comparing TTMs with AC-IVs (3). As already pointed out, our SSN analysis with a slightly less stringent alignment scores in the SSN merged the AC-IV-specific cluster with the *Sa*TTM cluster. Also, *Sa*TTM shared ∼29% identity with both TTMs and AC-IVs, so it is difficult to annotate these conserved enzymes solely on sequence similarity, as the function-specific features of CYTH enzymes have not been well defined. Our cluster analysis of the CYTH superfamily SSN identifies three key motifs that enable us to predict functions for all family members (Fig. 6, Table S1). According to our analysis, a true AC-IV requires a conserved HF-motif that extends the N-terminal signature ExExK-motif to HFxxxxExExK, a feature that is absent in all other *CYTH* enzymes. In the crystal structure of *Yp*CyaB the phenylalanine (F5) is involved in π-π stacking of the adenine base, while the histidine (H4) residue stacks to the phenylalanine via cation-π interaction, suggesting a key role in binding ATP in the correct orientation. When F5 residues were mutated in AC-IV family members their catalytic activity was drastically reduced (7). In addition, the adenine base stacks to the phenylalanine of the HF-motif within a relatively hydrophobic local environment, but no clear directional contacts like hydrogen bonds are made to the adenine amino group indicating that these AC-IVs act not strictly as adenylyl cyclases. The key HF-motif is absent in the sequence of all other CYTH proteins hence we infer that in the absence of an N-terminal HF-motif the CYTH enzyme function is either TTM or ThTPase.

**Figure 6.**
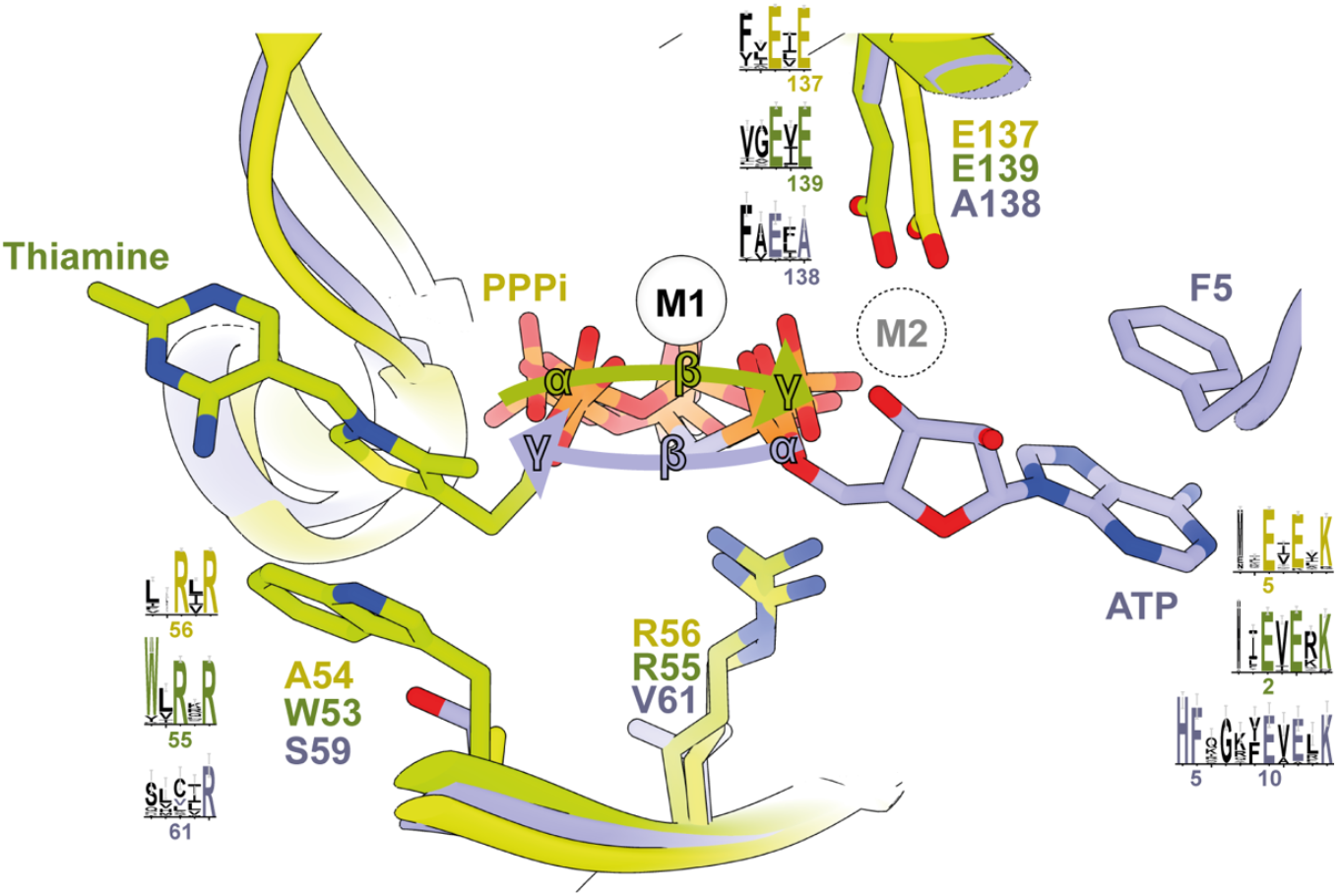
Key motifs for substrate orientation and specificity of PF01928 enzymes. Superposition of the a. Adenylyl cyclase class IV enzymes possess an exclusive, N-terminal HF-motif for nucleobase stacking interaction by the phenylalanine. This motif is located four residues before the CYTH hallmark motif ExExK. The second motif identifies ThTPases using an aromatic residue to stack the organic part of the thiamine-triphosphate using the WxRxR motif. TTMs and AC-IV have small residue substitutes and the cyclases further exchanged the otherwise conserved centered arginine to a small, neutral residue, which is not sterically hindering correct ribose placement. The third, acidic ExE-motif is responsible for correct metal coordination. The first glutamate is conserved throughout the while family, as it is the central residue for metal 1 coordination, while the latter glutamate establishes the interaction to metal 2, which is necessary for the catalytic mechanism in TTMs and ThTPases. This glutamate is strictly replaced by an alanine in the cyclase cluster 10, suggesting the inability of metal 2 coordination.

On the other hand, the W/YxRxR motif defines the ThTPases, as this subfamily has a conserved aromatic residue (W53 in case of mouse ThTPase) that is capable of stacking the aromatic thiazole of thiamine. Mutation of this residue reduced the plasticity of the ThTPase (6). Interestingly, we find this motif only in one other subfamily (cluster 7) that contains a bidomain class harboring enzymes of the plant TTM1/2 type, which are “TTMs” with a strong substrate preference for pyrophosphate (18). All other clusters have small amino acid substitution at this position (A54 for *Sa*TTM and S59 for *Yp*CyaB). Two arginine residues (R55 and R57 for mouse ThTPase) are required for interaction with the β- and γ-phosphates of ThTP. In AC-IVs, the first arginine residue in this motif is substituted by a small amino acid (V61 in case of *Yp*CyaB), mostly cysteine, valine or leucine (Fig. S6). Hence, it can be hypothesized that the conserved arginine, which interacts with the γ-phosphate in TTM/ThTPases, is a gate keeper against NTP binding in the cyclase orientation, as the guanidinium group otherwise sterically clashes with the ribose moiety (Fig. 6). The third characteristic feature is an ExE motif on strand β8 close to the C-terminus of CYTH proteins (E135 and E137 for *Sa*TTM). These conserved glutamate (E) residues are required for metal binding. In TTM/ThTPases, the second glutamate residue is required for metal 2 binding, which is strictly substituted to alanine in case of AC-IVs. Although this contradicts the generic binding of a second metal ion, a two-metal AC-IV mechanism was suggested before for this subfamily (7). Taken together, this combined analysis provides a basis to delineate proper functional annotation of all CYTH domain containing orthologs.

### Biological role of CYTH

The CYTH superfamily proteins exist in all kingdom of life and can be traced back to the last common ancestor, but the physiological function of this protein family has largely remained elusive. It has been proposed that members of CYTH superfamily might be involved in the interface of metabolism of phosphoryl compounds and nucleotides. Deletion mutants of the founding member of CYTH protein superfamily showed no significant cellular defects (21). In addition, the unusual enzymatic properties, notably the basic pH and high temperature optima indicate that CYTH enzymes might hardly play a physiological role under normal growth conditions.

So far, the best characterized CYTH proteins from a functional viewpoint derive from plants. The Arabidopsis TTM3 (*At*TTM3) knockout plant showed an anomalous root phenotype, indicating a potential role in root development (22). Although it has been established that plant TTMs act as a polyphosphate hydrolase, recent reports suggest that *Brachypodium distachyon* and *Hippeastrum x hybridum* TTM proteins might be able to produce a low amount of cAMP under laboratory conditions (23, 24). In our enzyme function-based cluster analysis (Fig. 1) we identified all of the so far characterized plant TTMs were present in the same cluster (cluster 9). As stated above, *At*TTM1/2 from cluster 7 are different from classical TTMs, as they lack the signature motifs for triphosphate binding and γ-phosphate hydrolysis (ExExK and ExE, see Table S1). This cluster still harbors the ThTPase typical W/YxRxR motif that is involved in substrate binding from the metal-opposing site, as well as the described DxY motif required for α /β-water placement (Fig. 5 and S8). This goes along with the observation that *At*TTM2 exhibits pyrophosphatase activity *in vitro*. Upon their deletion, plants became hypersensitive to both virulent and avirulent pathogens and accumulate elevated levels of sialic acid upon infection (25). On the other hand, *At*TTM1 that holds a similar enzyme activity as *At*TTM2 was shown to be rather involved in leaf senescence (18). These contrasting findings are food for future studies on the regulatory and metabolic roles of plant TTMs.

The strictly cAMP producing CYTH enzymes, the AC-IVs, form only a small cluster in the whole CYTH sequence similarity network (Fig. 1 cluster 10). The biological role of AC-IV is yet to be established. The seminal work discovering AC-IV in *Aeromonas hydrophila* showed that the *cyaB* gene is not expressed under standard growth conditions. However, it can complement the function of another adenylyl cyclase, *cyaA*, when overproduced in a *cyaA* deletion strain. The *cyaB* mutant strain had no visible defects and behaved similar to the wild type strain. Cellular cAMP levels were also noted to be unaltered upon deletion under normal growth condition (21). Interestingly, in our SSN analysis we identified that the huge majority of organisms that harbor an AC-IV are (potential) pathogens. Hence it is conceivable to envision that these AC-IVs might be repressed during normal growth and play hence no function on cellular cAMP production, but participate during pathogenesis. AC-IV enzymes possess no similarities with other bacterial toxins and are devoid of any signal sequence (21). Clearly, future studies will be required to address how expression of AC-IV enzymes is repressed under normal growth condition and what is their precise role during host infection and pathogen maintenance.

In this study we showed that the CYTH enzyme in crenarchaeote *Sulfolobus acidocaldarius* previously annotated as adenylyl cyclase is actually a phosphohydrolase of the TTM family. *Sa*TTM prefers inorganic triphosphate substrate over NTPs. Inorganic polyphosphates are considered as energy source readily available for converting di or mono phosphorylated nucleotides to more accessible tri-phosphorylated ones (like ATP), as metal chelating polymers or as buffers for cellular pH homeostasis (26). In bacteria, polyphosphates are formed by PolyP kinase (PPK), whose deletion mutants are defective in biofilm formation, quorum sensing, motility and other virulent properties (27). Electron dense amorphous PolyP granules were noted in the archaeal counterpart (28–30). PolyP accumulation in archaea is directly linked to metal toxicity (31) and oxidative stress (32) and its production is dependent of inorganic phosphate availability in *Methanosarcina mazei* (33). The exopolyphosphatase (PPX) gene was identified in *Sulfolobales*, while the polyP kinase is yet to be identified (34). In a recent study the overproduced *ppx* gene in *Sulfolobus acidocaldarius* led to a mutant strain lacking PolyP which was impaired in biofilm formation, motility, archaellum assembly and adhesion to glass surfaces (35). Among extremophiles polyP might play a crucial role in environmental fitness (34). Although the concentration of triphosphates is not known in *S. acidocaldarius* but considering our present finding and previous reports, we envision that TTM in archaea might be directly involved in the polyP metabolism and will indirectly take part in PolyP mediated regulatory processes and oxidative stress response.

## Experimental procedures

### Protein over production and purification

*Sa*TTM (*Saci_*0718) ORF amplified from Sulfolobus acidocaldarius DSM 639 genome having an *Nco*I and *Bam*HI restriction enzyme sites was cloned into *Nco*I/*Bam*HI digested T7-RNA polymerase based expression plasmid pSA4 (36) in order to produce a pSVA1028 that upon induction with IPTG produce C-terminally His6 fusion of *Sa*TTM. The site directed mutants were produced using ‘around-the-horn’ PCR of the pSVA1028. The correct DNA sequences were verified by sequencing.

*Escherichia coli* BL21-Gold (DE3) was used for heterologous overproduction of C-terminal His_6_ fused *Sa*TTM. Cells were grown in lysogeny broth supplemented with 100 µg/ml ampicillin at 37 °C until an OD_600_ of 0.5. The overexpression of SaTTM was induced by using 0.75 mM IPTG at 18 °C overnight. Cells from 2 L culture were harvested (4000 rpm for 20 min), resuspended in 30 ml lysis buffer (200 mM NaCl, 20 mM HEPES, pH 8.0) and subsequently lysed using French Press. The lysate was incubated at 70 °C for 10 min to reduce *E. coli* contaminants and the heat stable cell free lysate was collected using 18000 rpm for 20 min. The supernatant was loaded on 5 ml Ni-NTA column (Protino™) and washed with 6 CV lysis buffer before eluting the protein with 4 CV elution buffer (lysis buffer with 500 mM imidazole). The elution was concentrated and further purified by size exclusion chromatography using a HiLoad^®^ 26/600 Superdex^®^ 200 PG (GE Healthcare) column equilibrated with SEC-buffer (200 mM NaCl, 50 mM Tris, pH 9.0). SaTTM protein purity was monitored using a Coomassie stained 15% SDS-PAGE. The oligomerization of SaTTM was checked on an analytical Superdex^®^ 200 10/300 GL column and using SEC buffer system. Pure protein was concentrated and stored at −80 °C until activity assays or crystallization.

### Crystallization of *Sa*TTM

To obtain *Sa*TTM crystals 10 mg/ml protein in SEC-buffer was mixed with the respective precipitant in an equal volume ratio. Crystals were grown at 4°C using sitting-drop vapour diffusion. For the ATP state, 10 mM CaCl_2_ and 5 mM ATP were added to the protein solution. The best diffracting crystal appeared in condition JCSG Core IE7 (0.2 M Calcium acetate, 0.1 M MES pH 6.0, 20% (w/v) PEG 8000). The relaxed state of *Sa*TTM (PDB: 7NS8) originated from the ATP screen in a precipitant condition with 0.2 M Lithium sulphate, 0.1 M CAPS pH 10.5, 2 M ammonium sulphate. High concentration of sulphate in the precipitation condition competed out the phosphate substrate from the active site. The Michaelis complex was obtained from a SaTTM mixed with 10 mM CaCl_2_ (or 10mM MgCl_2_), 0.5 mM MgCl_2_, and 5 mM of hexa-polyphosphate, P_6_i (or tri-polyphosphate, PPPi). Both, PPPi+Ca^2+^ and PPPi+Mg^2+^, crystallized in 0.2 M NaCl, 0.1 M Na/KPO_4_ pH 6.2, 20% (w/v) PEG 1000. Note, the hexa-polyphosphate was highly impure as detected by malachite green assay (data not shown). The product (PPi) inhibited SaTTM in presence of 10mM CaCl_2_ and 5mM PPi was crystallized in 0.17 M ammonium acetate, 0.085 M sodium acetate pH 4.6, 25.5% (w/v) PEG 4000, 15% (v/v) glycerol. Crystals were harvested at 4°C as temperature seems to have significant effect on crystallization of SaTTM complexes. Crystals were cryoprotected in mother liquor supplemented with 30% glycerol except the PPPi-Mg^2+^ complex and subsequently flash frozen using liquid nitrogen for data collection.

### Structure solution

X-ray data for different states of *Sa*TTM were collected from single crystals at the ESRF (Grenoble, France). Crystallography data were processed using iMOSFLM in the CCP4i2 package (37). Initial phases for *Sa*TTM-ATP-Ca^2+^ were obtained using a *SWISS-model* (38) of *Sa*TTM, based on PDB entry 2EEN (identity 42%) as a search model for molecular replacement in Phaser-MR (39). Atomic models of the other crystallized states of *Sa*TTM were obtained by using the structure of *Sa*TTM in complex with ATP and calcium as a search model for Phaser-MR. Iterative model building and refinements was performed with COOT and Phenix.refine (40, 41). Data collection and refinement statistics are summarized in Table S2. The resulting coordinates were deposited at the protein data bank under accession numbers 7NS8, 7NS9, 7NSA, 7NSF, and 7NSD. The structural analysis and figure generation was performed with UCSF Chimera (42).

### Activity assay

Triphosphatase activity of *Sa*TTM was analysed with the Malachite Green Phosphate Assay Kit (MAK307-1KT) from Sigma Aldrich according to manufacturer’s protocol. Reactions were quenched by mixing 80 µl assay mix into 20 µl working reagent using 96 well PS F-bottom microplates (Greiner). After colour development for 30 min at room temperature, the absorbance was measured at 620 nm using a plate reader (photospectrometer Infinite 200, Tecan). The results were analysed with Microsoft Excel and GraphPad Prism. The pH optimum of *Sa*TTM was determined by measuring the amount of reaction product (Pi release) from 5 nM *Sa*TTM, 2 mM MgCl2, and 0.1 mM PPPi buffered in 50 mM Tris and 200 mM NaCl at varying pH values (6.1-9.1) after 5 min incubation. To determine the preferred metal cofactor, 5 nM *Sa*TTM, 2 mM metal ion (Mg^2+,^ Ca^2+^, Mn^2+,^ Zn^2+,^ Ni^2+)^ and 0.1 mM PPPi were incubated at 75°C for 5 min buffered in 50 mM Tris and 200 mM NaCl at pH 8.6. The preferred substrate was analysed by using the same parameters for 10 min with Mg^2+^ as a cofactor and varying substrates (PPPi, ATP, GTP, AP4A, AP5A) at 0.1 mM concentration. Mutant activities were determined accordingly with 5 min incubation. In order to test the product inhibition, as well as the ability of Ca^2+^ to inhibit *Sa*TTM, a titration series of 1 nM to 10 mM inhibitor (PPi or Ca^2+)^ was performed by measuring their influence on the product formation of 5 nM *Sa*TTM, 2 mM MgCl_2_, and 0.1 mM PPPi buffered in 50 mM Tris and 200 mM NaCl at pH 8.6 incubated for 5 and 10 min at 75 °C.

To analyse the enzyme kinetics of *Sa*TTM phosphatase activity, 5 nM enzyme was incubated together with 2 mM MgCl_2_ and varying PPPi concentrations (0.005 to 0.5 mM) incubated in 50 mM Tris and 200 mM NaCl at pH 8.6 and 75 °C. To obtain kinetic parameters, the velocities at different substrate concentrations were determined by measuring the Pi formation at different time points and the resulting linear regressions of four measurements were used for the Michaelis-Menten analysis.

To check products on HPLC, the reactions were diluted 5-fold with double-distilled water and the enzyme denatured by adding 100 µl chloroform followed by 15 s of vigorous shaking, 15 s heat denaturation at 95 °C and snap freezing in liquid nitrogen. The samples were thawed, the two phases separated by centrifugation (17,300 *x g*, 10 min, 4 °C) and 10 µl of the aqueous phase containing the nucleotides subjected to HPLC analysis. Nucleotides were separated with an anion exchange column (Metrosep A Supp 5 – 150/4.0) with 100 mM (NH_4_)_2_CO_3_ Ph 9.25 at a flow rate of 0.7 ml/min as eluent and detected at 260 nm wavelength in agreement with standards.

For the activity analysis of *Sa*TTM towards P_4_, 21 μM *Sa*TTM were incubated in a total volume of 150 μl together with 15 mM Tris pH 7.5 and 4 mM MgCl2 for 3 min at 75 °C. The reaction was initiated by the addition of 0.5 mM P_4_ and terminated after 10 min through incubation on ice for 10 min, followed by centrifugation in a spin filter to remove the enzyme (4 °C, 35 min, 10.0 x1000 rcf, Amicon-Ultra 0.5, 10 kDa). The flow through was analysed by ionic exchange chromatography using a Thermo Scientific Dionex™ ICS-5000+ equipped with a conductivity detector and an EGC500 eluent generator. 10 µl of the flow through were injected followed by elution with a flow rate of 0.25 ml/min and a KOH gradient starting from 5 mM KOH. The eluent was raised to 12 mM within 5 min, followed by an increase to 20 mM until 10 min and 70 mM within 15 min. KOH concentration was maintained at 70 mM until 26 min, subsequently lowered to 5 mM at 26.1 min followed by equilibration until termination of the run at 27 min.

### Sequence similarity network

The protein family PF01928 (CYTH) was used as input for the Enzyme Similarity Tool provided by the Enzyme Function Initiative (14) to generate the sequence-similarity network (SSN). Using the domain boundaries and an E-value of 10^−5^, the initial network was generated and subsequently modified by applying higher stringent cut offs (10^−15^, 10^−20^) as alignment scores for the final SSNs. Each node represents proteins with >40% identity. The resulting networks were subjected to the recently implemented cluster analysis for automatic cluster-WEBLOGO generation and the final networks were analysed with Cytoscape 3.5.1 (43, 44).

## Data availability

The atomic coordinates of the crystal structures of *Sa*TTM, *Sa*TTM/PPPi/Ca^2+^, *Sa*TTM/PPi/Ca^2+^, *Sa*TTM/PPPi/Mg^2+^, and *Sa*TTM/ATP/Ca^2+^ have been deposited in the Protein Data Bank (PDB ID codes 7NS8, 7NS9, 7NSA, 7NSF, and 7NSD, respectively).

## Acknowledgments

We thank Dr. Xing Ye for his guidance and advises and the group of Prof. Henning J. Jessen (Institute of Organic Chemistry, Albert-Ludwigs-Universität Freiburg) for substrate and IC system supply. We also thank Ralf Pöschke (MarXtal facility, Marburg) for technical support during crystallization and the staff of beamlines ID23-1, ID29, and ID30B at the European Synchrotron Radiation Facility (Grenoble, France) for support with data collection. Further, we thank Dr. Lena Hoffmann for cloning the initial *Sa*TTM vector, Julian Grützner and Abbas Sajediabkenar for their help during their internships, and Dr. Wieland Steinchen for assistance with HPLC measurements.

## Author contributions

MSV, RRN, MKFM and AB performed experiments. MSV, LOE and AB planned experiments, analyzed data, prepared initial draft, and finalized the manuscript draft. LOE and SVA revised the manuscript.

## Funding and additional information

This work was supported by DFG grant SFB 987 (Collaborative Research Center 987) and Volkswagen Foundation (Az96727) to LOE, MSV and SVA. AB is supported by DFG fellowship (666472).

## Supplementary to

## Abbreviations

*Pi*: orthophosphate
*PPi*: pyrophosphate
*PPPi*: triphosphate
*CYTH*: CyaB-Thiamine triphosphatase

## Supplementary figures

**Figure S1.**
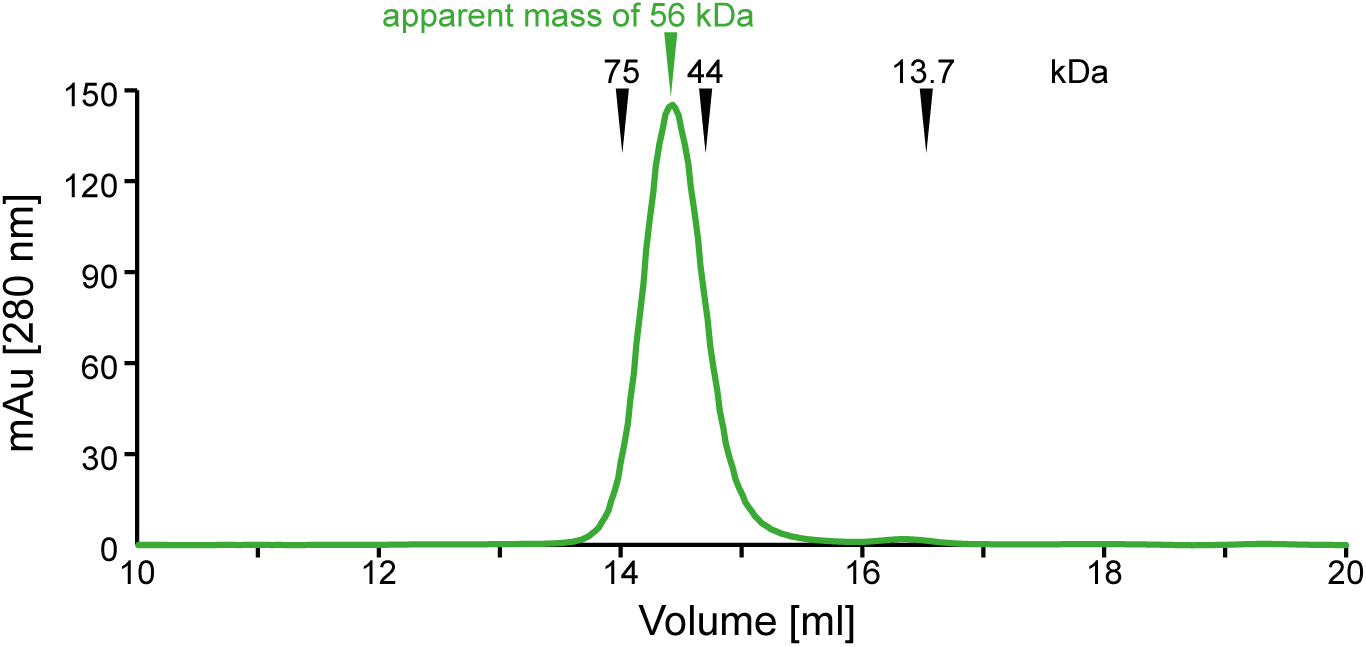
Calibrated SEC run of *Sa*TTM on Biorad SEC650 column. Conalbumin (75 kDa), ovalbumin (44 kDa), and ribonuclease A (13.7 kDa) were used as mass standards. The calculated molecular mass of the *Sa*TTM monomer is 23.65 kDa, its SEC elution corresponds to a dimer in solution.

**Figure S2.**
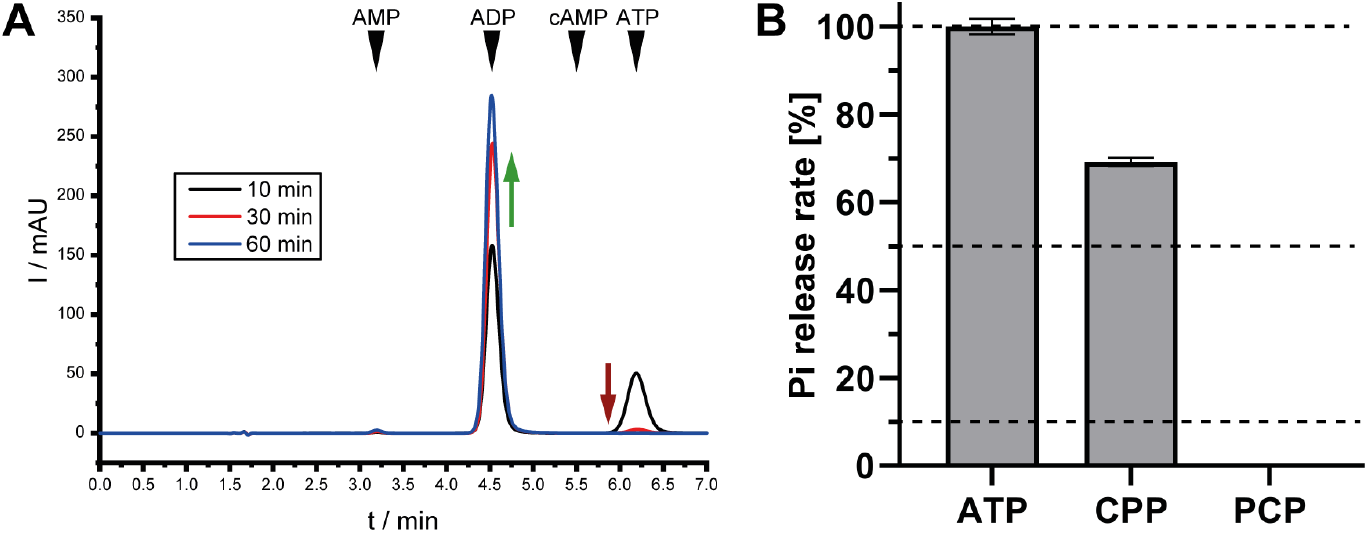
*Sa*TTM converts ATP to ADP by releasing γ-phosphate. **A** Chromatograms of the time course of *Sa*TTM hydrolyzing ATP to ADP is shown. As indicated by standards, no cAMP is produced. Arrows indicate the decrease (red) of substrate increase (green) of product over time. **B** *Sa*TTM can utilize ATP and the non-hydrolysable ATP analog AMPCPP, but not AMPPCP, as a substrate.

**Figure S3.**
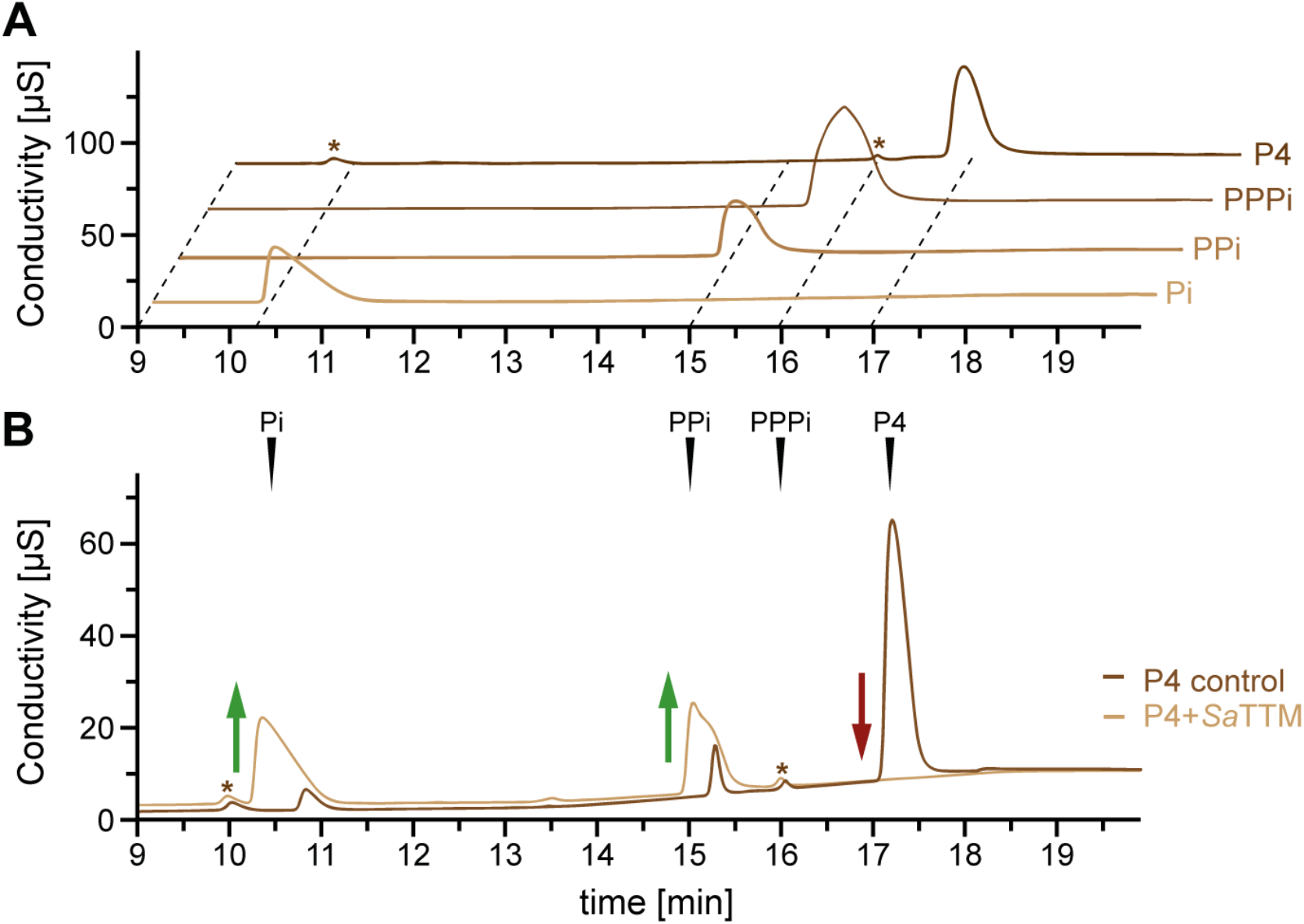
*Sa*TTM can hydrolyze tetraphosphate. **A** Chromatograms of standards are shown for phosphate (Pi), pyrophosphate (PPi), triphosphate (PPPi), and tetraphosphate (P4). **B** Chromatograms of P4 after incubation alone (brown) and together with *Sa*TTM (ocher) show a complete turnover of P4 as indicated by the red arrow, while the products Pi and PPi increase (green arrows). The two unidentified peaks indicated by asterisks appear throughout the ion exchange chromatography run of P4 without being affected by the sample.

**Figure S4.**
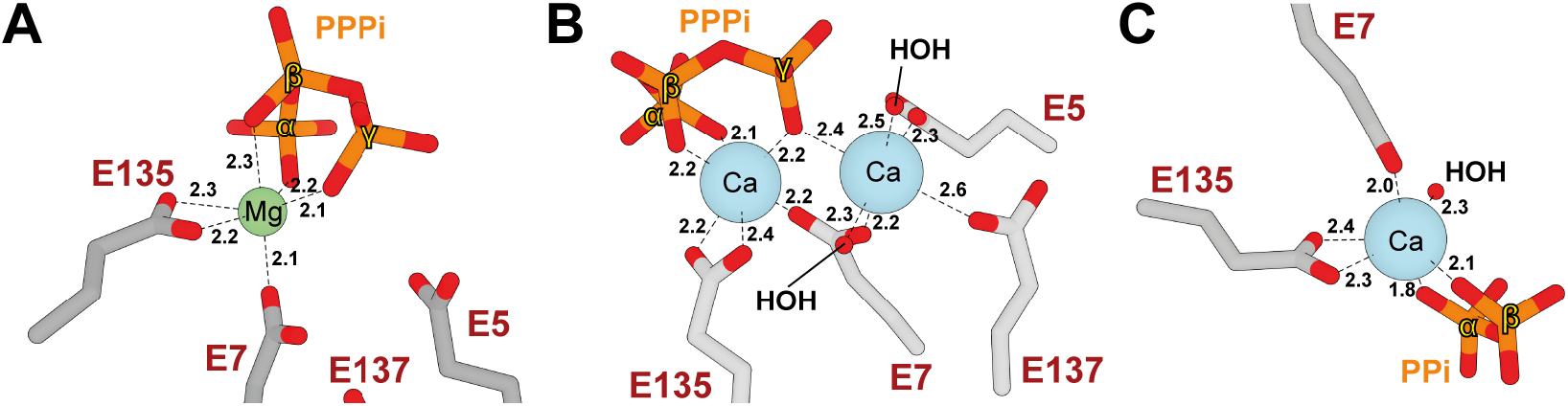
Distorted pseudo-octahedral metal ion coordination in *Sa*TTM. The metal coordination of *Sa*TTM PPPi•Mg^2+^ (**A**), PPPi•Ca^2+^ (**B**), and Ppi•Ca^2+^ (**C**) is shown indicated by the dashed lines and the respective distances shown in Angstrom. α -, β-, and γ-positions of phosphates are indicated, so are the residues shown as grey sticks. Oxygens are colored in red, phosphorus in orange, magnesium as a green sphere and calcium as blue sphere. Waters are red spheres as indicated.

**Figure S5.**
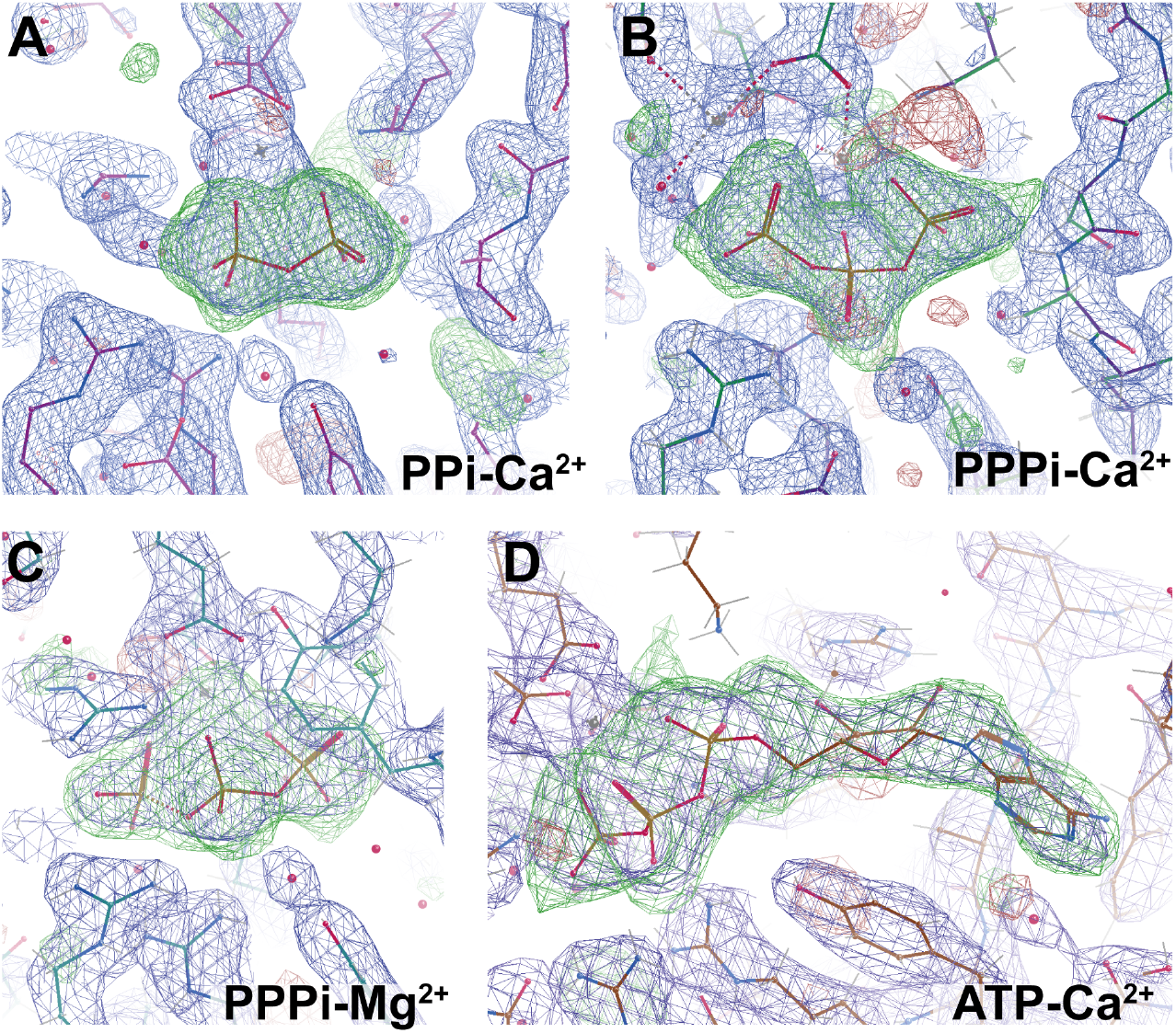
OMIT maps for the different catalytic states of *Sa*TTM. OMIT maps of the respective ligands contoured at σ-level 3.0 are shown in the context of SIGMAA-weighted 2m*F*o-D*F*c maps (blue mesh, contouring level: 1.5) of nearby residues.

**Figure S6.**
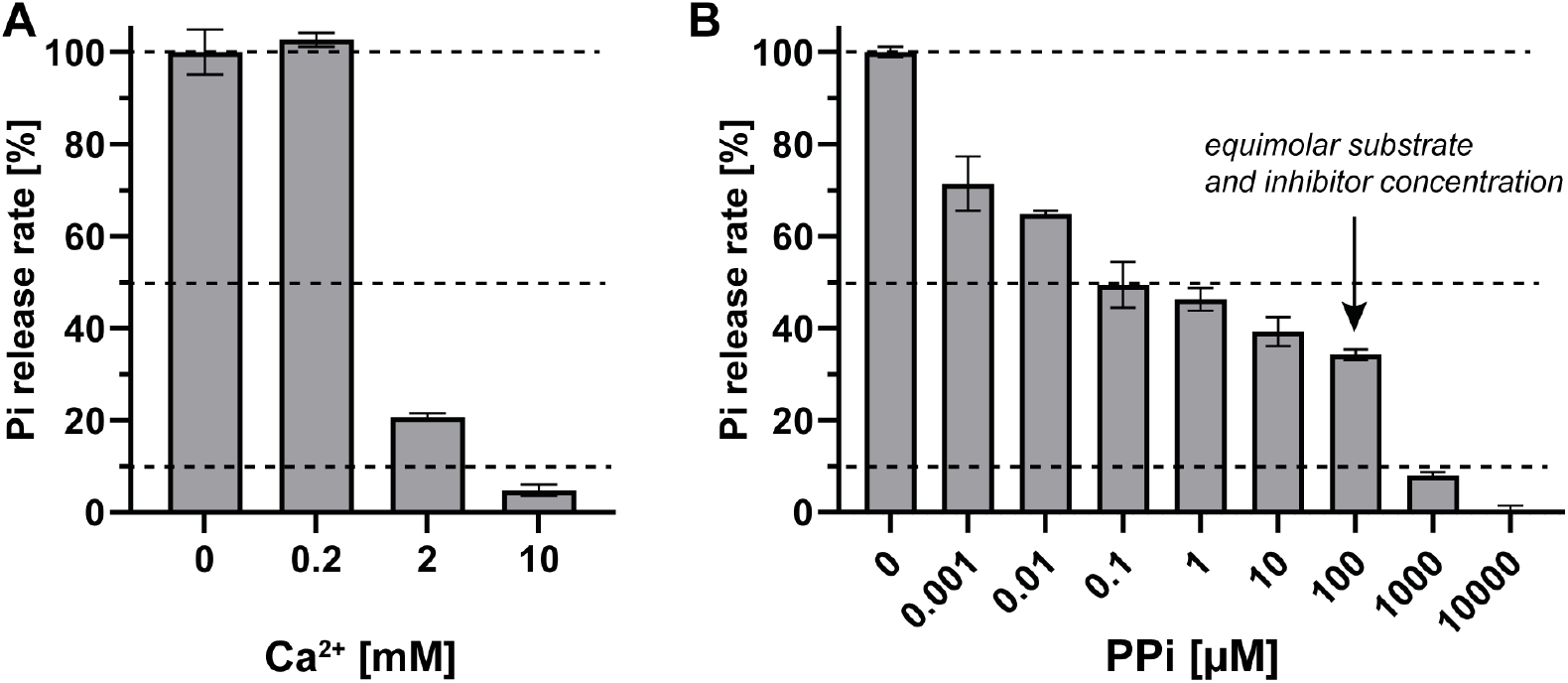
Titrations with inhibitory calcium and the product PPi. **A** The inhibiting effect of calcium on the triphosphatase activity of *Sa*TTM is shown. **B** The presence of the product PPi has an inhibitory effect on the reaction behavior of *Sa*TTM.

**Figure S7.**
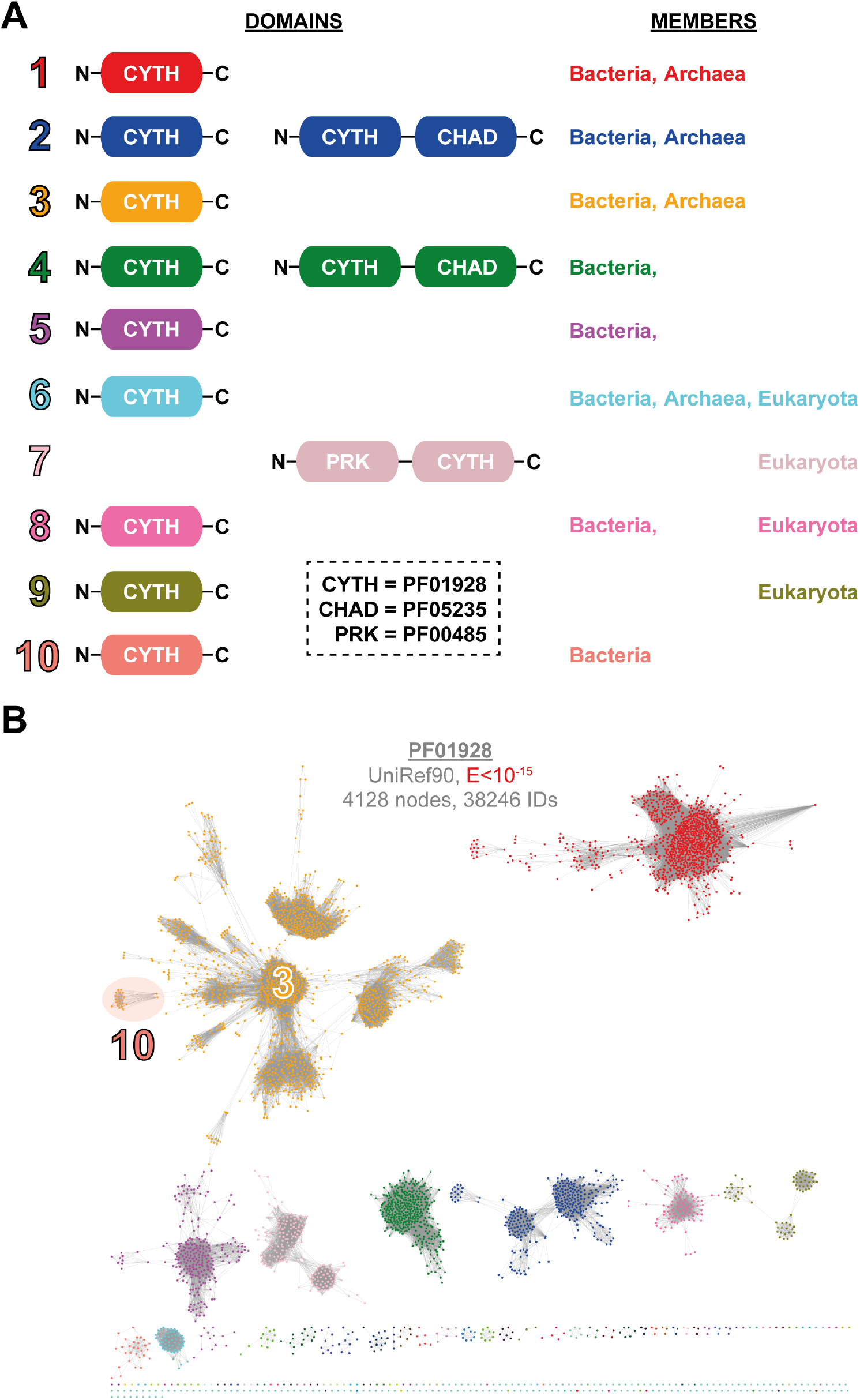
Domain architectures of PF01928 family and its SSN with lower stringency. **A** The prominent domain architectures present in the ten described clusters of the PF01928 family are shown with their appearance among the three kingdoms. The Pfam IDs are noted in the dashed box. **B** The PF01928 SSN with an alignment score of E<10^−15^ is presented. Cluster 10 from the SSN in Fig. 1 is now member of cluster 3 (highlighted in transparent red).

**Figure S8.**
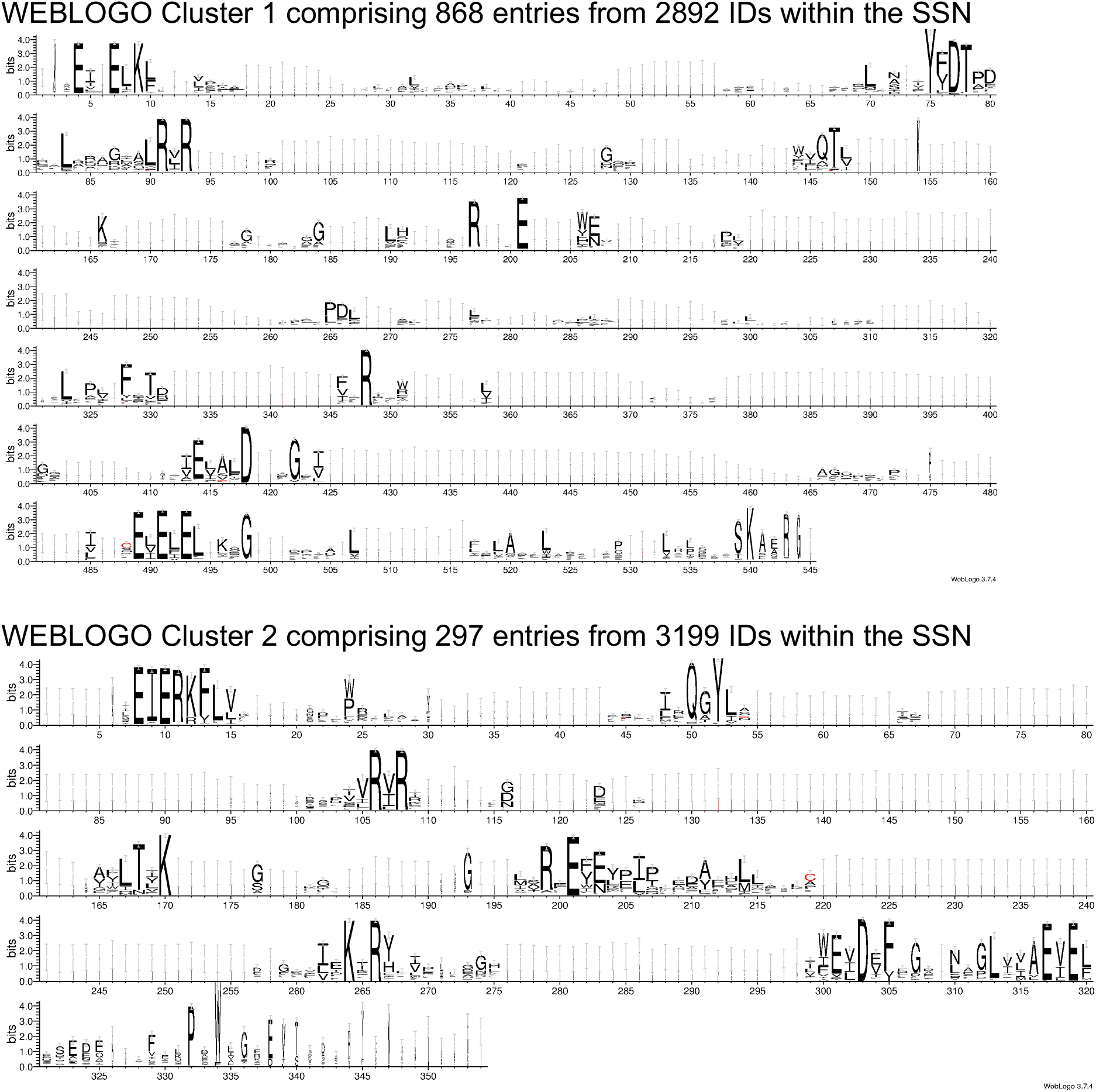
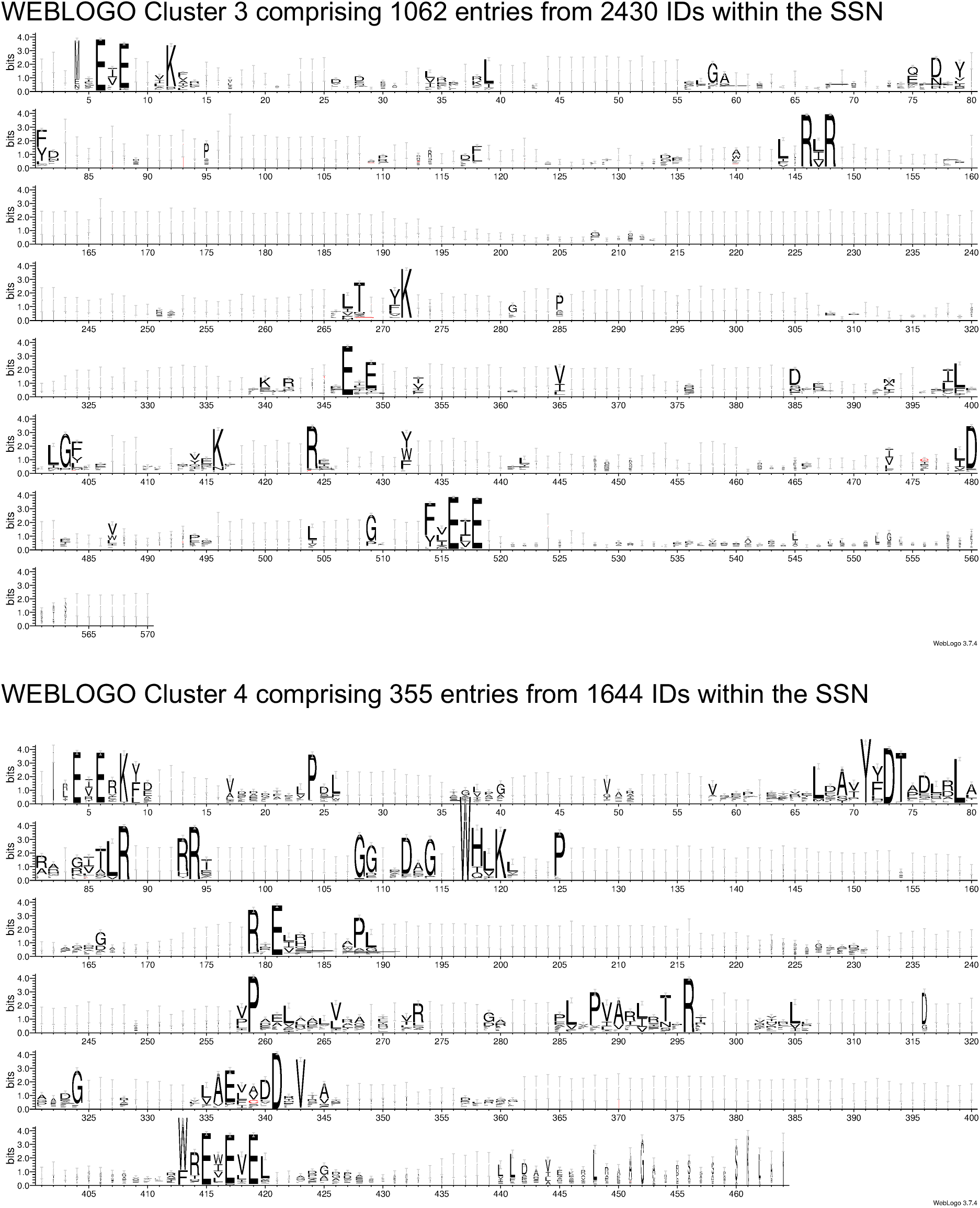
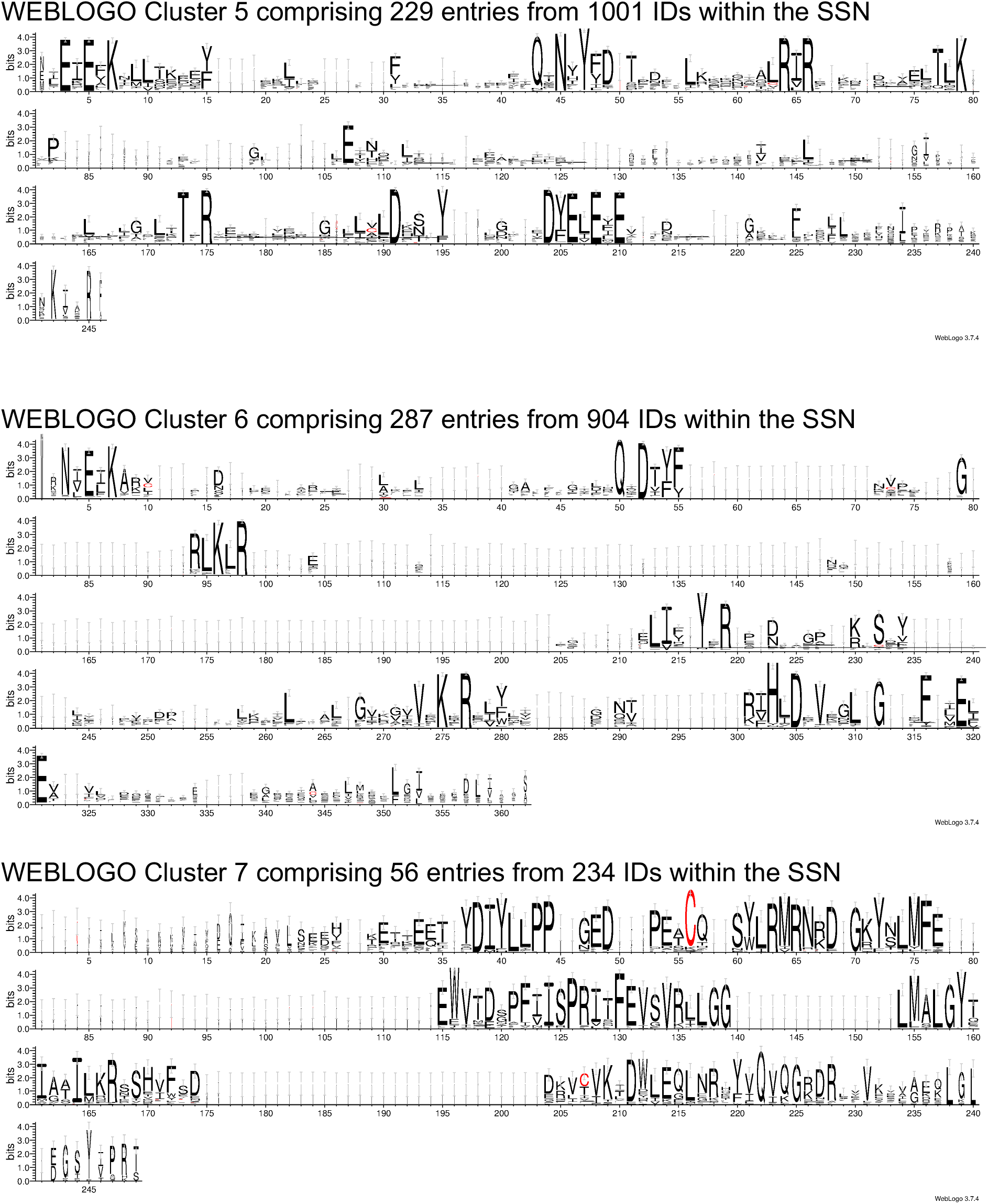
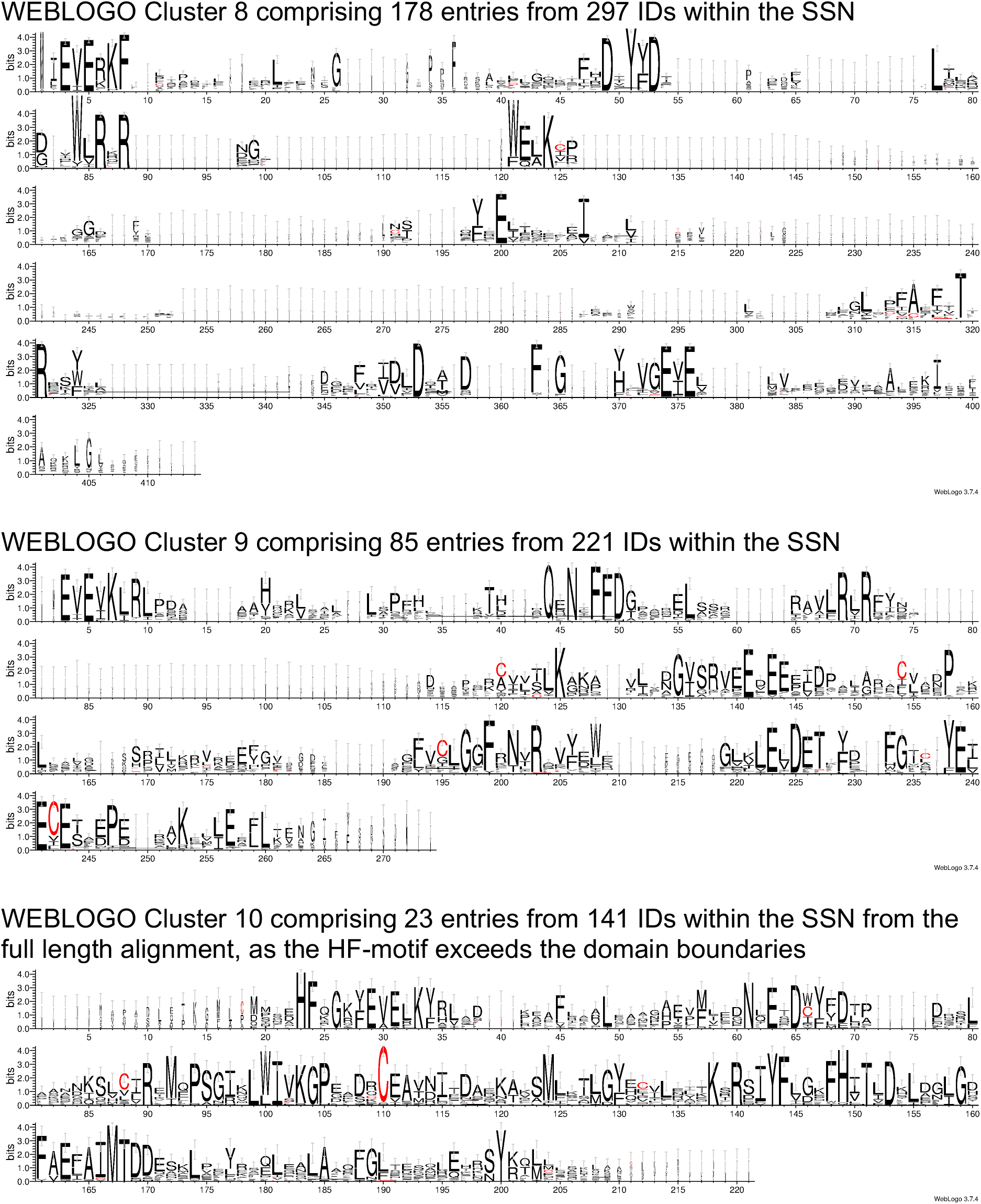
WEBLOGOS of PF01928 clusters 1-10.

## Supplementary tables

**Table S1:**
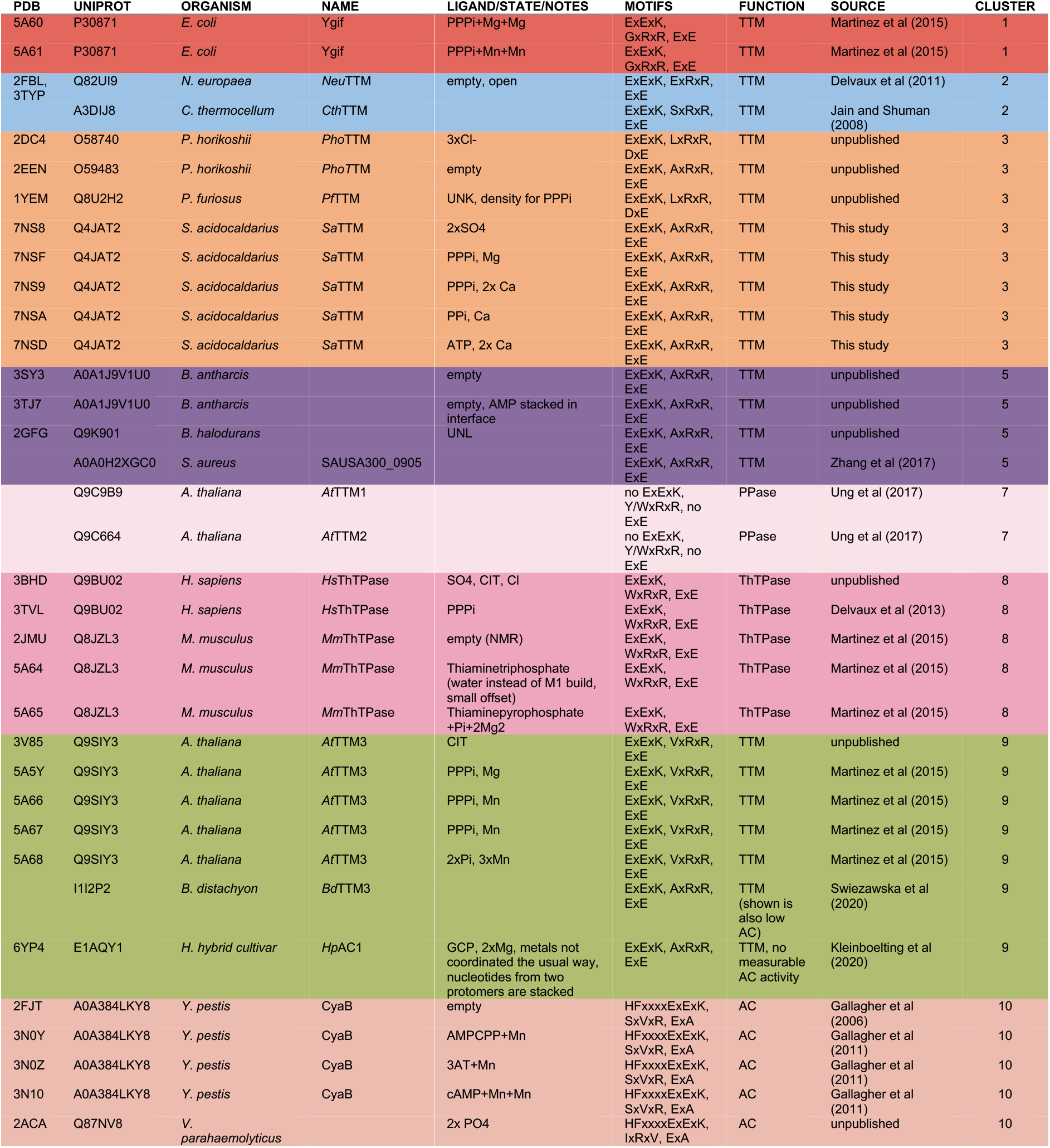
Overview of characterized PF01928 enzymes. Coloring according to SSN.

**Table S2:** Data collection and refinement statistics

| SaTTM | relaxed/SO<sub>4</sub> | PPPi/Ca<sup>2+</sup> | PPPi/Mg<sup>2+</sup> | PPi/Ca<sup>2+</sup> | ATP/Ca<sup>2+</sup> |
| --- | --- | --- | --- | --- | --- |
| *PDB code* | 7NS8 | 7NS9 | 7NSF | 7NSA | 7NSD |
| *Data collection* |  |  |  |  |  |
| Synchrotron beamline | ESRF ID29 | ESRF ID29 | ESRF ID23-1 | ESRF ID30B | ESRF ID29 |
| Wavelength (Å) | 0.979 | 0.979 | 0.972 | 0.976 | 0.979 |
| Resolution range (Å) | 44.4-2.3 (2.38-2.30) | 46.0-1.75 (1.81-1.75) | 43.2-2.0 (2.07-2.0) | 46.0-1.95 (2.02-1.95) | 45.8-2.2 (2.27-2.19) |
| Space group | *P*4<sub>1</sub>2<sub>1</sub>2 | *P*4<sub>1</sub>2<sub>1</sub>2 | *P*4<sub>1</sub>2<sub>1</sub>2 | *P*4<sub>1</sub>2<sub>1</sub>2 | *P*4<sub>1</sub>2<sub>1</sub>2 |
| Unit cell (Å, °) | 57.66, 57.66, 139.15,<br>90, 90, 90 | 60.67, 60.67, 141.09,<br>90, 90, 90 | 61.0, 61.03, 136.62,<br>90, 90, 90 | 60.55, 60.55, 141.63,<br>90, 90, 90 | 60.67, 60.67, 139.35,<br>90, 90, 90 |
| Total reflections | 22129 (2148) | 51470 (5143) | 36416 (3558) | 38954 (3885) | 28144 (2738) |
| Unique reflections | 11068 (1074) | 25885 (2581) | 18208 (1778) | 19769 (1955) | 14072 (1367) |
| Multiplicity | 2.0 (2.0) | 2.0 (2.0) | 2.0 (2.0) | 2.0 (2.0) | 2.0 (2.0) |
| Completeness (%) | 99.95 (100.00) | 94.31 (96.81) | 98.92 (99.94) | 98.87 (99.69) | 99.95 (99.78) |
| Mean *I*/$\sigma$(*I*) | 29.77 (15.39) | 11.44 (2.12) | 17.55 (2.43) | 16.60 (2.65) | 17.15 (4.12) |
| Wilson B-factor (Å<sup>2</sup>) | 34.52 | 32.86 | 43.90 | 37.37 | 48.39 |
| *R*<sub>merge</sub> | 0.01 (0.04) | 0.02 (0.30) | 0.01 (0.26) | 0.02 (0.27) | 0.01 (0.17) |
| *R*<sub>meas</sub> | 0.02 (0.06) | 0.03 (0.41) | 0.02 (0.37) | 0.03 (0.38) | 0.02 (0.24) |
| CC1/2 | 1 (0.99) | 1 (0.35) | 1 (0.88) | 1 (0.73) | 1 (0.86) |
| *Refinement statistics* |  |  |  |  |  |
| Resolution range (Å) | 44.4-2.3 | 46.0-1.75 | 43.2-2.0 | 46.0-1.95 | 45.8-2.2 |
| *R*<sub>work</sub>/*R*<sub>free</sub> (%) | 22.6/25.7 | 18.9/23.5 | 20.7/23.9 | 22.5/24.6 | 22.4/25.3 |
| Average B-factor (Å<sup>2</sup>) | 41.38 | 51.48 | 59.86 | 47.2 | 63.42 |
| No. of atoms | 1476 | 1641 | 1607 | 1619 | 1572 |
| macromolecules | 1341 | 1477 | 1479 | 1465 | 1455 |
| ligands | 24 | 26 | 14 | 27 | 37 |
| solvent | 111 | 138 | 114 | 127 | 80 |
| R.m.s.d., (bonds, Å) | 0.010 | 0.007 | 0.008 | 0.018 | 0.011 |
| R.m.s.d., (angles, °) | 1.15 | 0.73 | 1.09 | 1.93 | 1.48 |
| Ramachandran plot |  |  |  |  |  |
| favored (%) | 97.44 | 97.16 | 97.71 | 97.69 | 96.55 |
| allowed (%) | 2.56 | 2.84 | 2.29 | 2.31 | 3.45 |
| outliers (%) | 0.00 | 0.00 | 0.00 | 0.00 | 0.00 |
| Rotamer outliers (%) | 2.05 | 0.62 | 0.62 | 2.47 | 0.00 |
| Clashscore | 3.68 | 1.34 | 3.70 | 4.00 | 3.04 |

